# Structural dynamics insights into principles underlying the fitness of new broadly potent AAVs

**DOI:** 10.64898/2026.03.24.713814

**Authors:** Molly E. Johnson, Bilge E. Ozturk, Thomas H. Tugwell, Maryl Lambros, Haley N. Janowitz, Keevon Flohr, Morgan Sedorovitz, Laura Campello, Jane E. Hartung, Brett Hogle, Megan Gillespie, Hannah Schriever, Hamzah Aweidah, Janis Koester, Inga Clausen, Stefan Seeber, Franco Revelant, Richard Schreurs, Fabian Koechl, Paul A. Sieving, José-Alain Sahel, William R. Stauffer, Rui T. Peixoto, Sascha Fauser, Rob Lin, James F. Conway, Susana da Silva, Jacek Krol, Miguel Betegon, Leah C. Byrne

## Abstract

Adeno-associated virus (AAV) is a leading platform for gene therapy, but current clinical-stage vectors require high doses associated with adverse events. Engineering of AAVs has produced more efficient vectors, although the mechanism underlying these improvements often remains poorly understood, limiting further development and raising potential safety concerns. Here, we leveraged a new workflow for AAV engineering with single-cell resolution, called scAAVengr-Hunt, to create best-in-class AAVs for gene delivery. ATX002, the top-performing vector, demonstrates broad potency across species, including nonhuman primate, mouse, and human, as well as across retina and brain. To understand the mechanism underlying this broad potency, we performed molecular dynamics simulations comparing AAV variants spanning a range of fitness levels. Structural dynamics analysis revealed a bifunctional molecular mechanism that confers potency through increased affinity of the capsid to the AAV receptor and regulation of heparan sulfate binding. This work provides critical insights relating structural mechanism to the fitness of engineered AAVs and establishes rich new avenues for AAV engineering through the integration of sequence-level analysis with computational biophysics.

## Introduction

Adeno-associated virus (AAV) is a leading vector for gene therapies, with hundreds of clinical trials underway ^1,2^. However, current clinical-stage AAVs are inefficient, and must be injected at high doses that are associated with immune response and toxicity, reducing the efficacy of the gene therapy, and in some cases, leading to severe adverse events, including death in multiple clinical trials ^3,4^. To achieve success, gene therapy vectors must transduce tissues at a low dose while still delivering a therapeutic payload to cells ^5–7^. Engineering AAVs with efficient delivery to targeted tissues and cell types remains a critical hurdle to the broad application of safe gene therapies.

To create AAV vectors with high translational potential, we developed the scAAVengr-HUnT (<u>s</u>ingle-<u>c</u>ell <u>AAV eng</u>ineering with <u>h</u>igh <u>un</u>biased throughput) platform. The scAAVengr-HUnT workflow offers a quantitative, systematic, and high-throughput approach to develop optimized AAVs for gene therapy. Using a single-cell RNA-Seq-based process, mRNA expression levels of highly diverse capsid libraries are simultaneously and quantitatively assessed head-to-head, with single cell resolution, *in vivo*.

Here, we applied the scAAVengr-HUnT workflow to intravitreal (IVT) administration of AAVs working in nonhuman primate (NHP) retina. The retina is the light sensitive tissue lining the back of the eye. Most retinal degenerative diseases involve photoreceptor cells in the outermost retina. However, passage of AAV from the vitreous to the outer retina through structural barriers, including the inner limiting membrane (ILM) and cellular layers, remains a challenge^8^. IVT injections, where the needle is positioned in the fluid-filled vitreous cavity, are low-risk in-office procedures in ophthalmology and easier to perform than surgical subretinal injections, in which the needle penetrates and lifts the neural retina to deliver AAVs to outer retina. While IVT injections allow for pan-retinal transduction through diffusion of AAVs in the vitreous, subretinal injections target a limited area of the retina, and are associated with iatrogenic damage. Yet, IVT delivery remains inefficient, and subretinal injections are the most common method for gene therapy delivery to the outer retina^9^. While recent efforts have improved intravitreally-delivered AAVs, overall efficiency in large animal models has remained low ^10,11^.

Using scAAVengr-HUnT, we show the creation of a new tier of AAVs, including ATX002, that is 14X (1.16log10) more potent than the current clinical-stage engineered vector 7m8^12^, currently in use in multiple clinical trials for wet adult macular degeneration, retinitis pigmentosa, and optogenetics. ATX002 achieves strong expression across both central and peripheral retina. We also show that while ATX002 was optimized in the primate retina, it also outperforms 7m8 in mouse retina, demonstrating that ATX002’s properties translate across species. ATX002 additionally outperforms naturally occurring AAV controls in mouse and NHP CNS tissue following intra-cisterna magna (ICM) administration, demonstrating efficiency across tissues. Further, we use molecular dynamics (MD) to investigate the mechanism underlying improved fitness. Although considerable effort has been devoted to engineering novel AAVs with enhanced transduction efficiency, these efforts have lacked guidance from biophysical and mechanistic insights. Here, we present what is, to our knowledge, the first mechanistic investigation of engineered AAVs through structural dynamics to uncover the molecular features that confer improved fitness. Critically, these findings establish general principles for future AAV design, moving beyond sequence alone to incorporate capsid structural dynamics.

## Results

### scAAVengr-HUnT platform identifies highly potent AAV variants

We first generated a highly diverse library of AAV variants based on AAV2 (Fig. 1A). A trimer controlled 6mer insertion (TC6) in which every amino acid is represented at every insert residue, was introduced at position 588 of VP1, which is an exposed loop on the capsid that has been shown to interact with heparan sulfate (HS), a primary receptor for AAV2 (fig. S1A,B). The library transgene cassette contained a ubiquitous CMV promoter driving expression of eGFP, followed by a unique 20-bp barcode to track the performance of each AAV variant. Deep sequencing of plasmid and packaged AAV libraries provided capsid/barcode pairings.

**Fig. 1:**
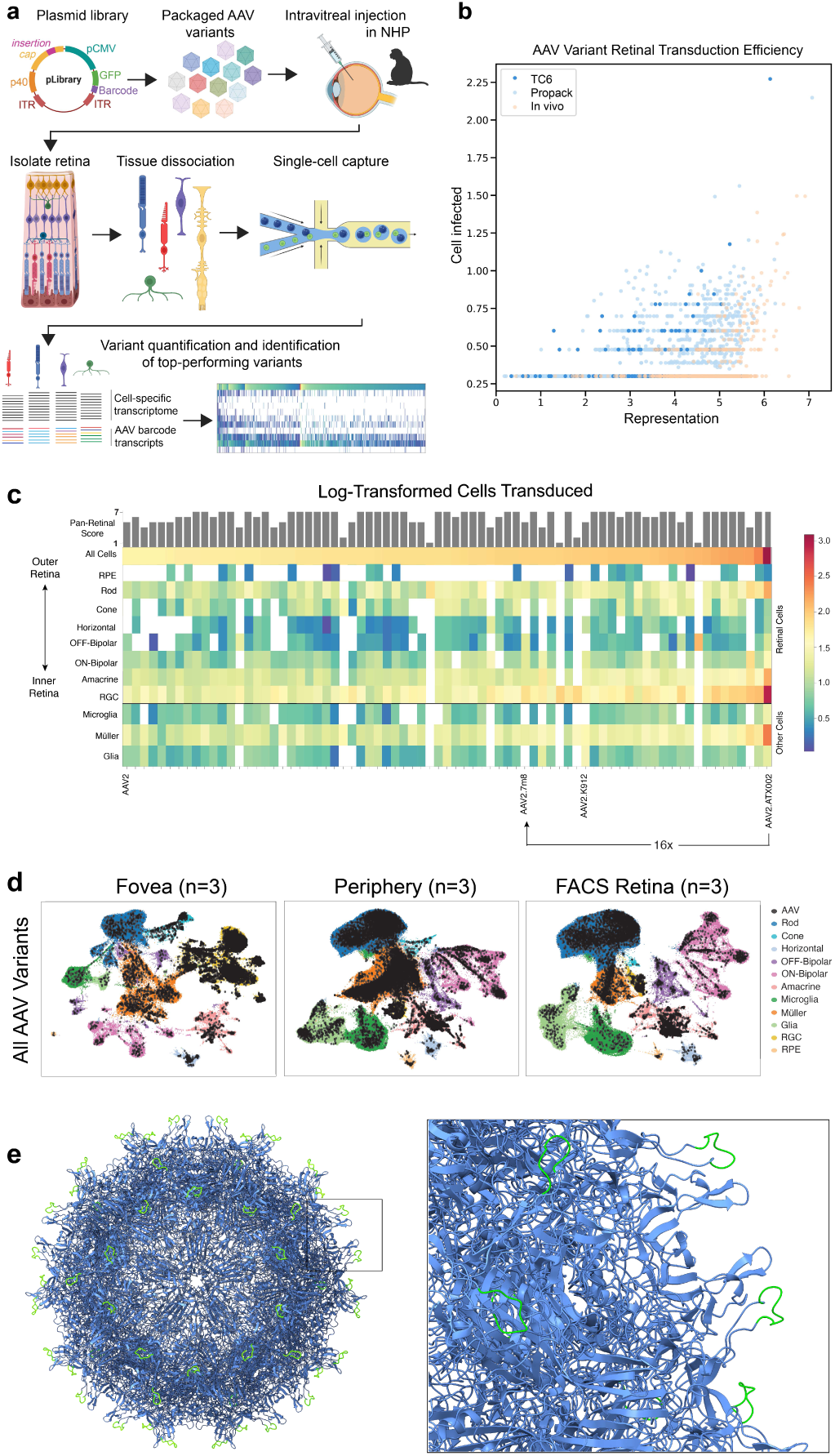
scAAVengr-HUnT platform and identification of top pan-retinal performer ATX002. **a,** Schematic of the scAAVengr-HUnT platform. Highly diverse capsid variant libraries are cloned, packaged and injected intravitreally. Single cell RNA-Seq processing and analysis is conducted to identify top performers. **b,** Scatter plot showing the distribution of retinal cells transduced vs. starting representation of AAV2-based capsid variants. The TC6 library is shown in dark blue, pro-pack library is in light blue, and *in vivo* enriched library is in light orange. **c,** Heatmap of top 200 AAV2-based variants tested alongside controls in a low-throughput scAAVengr screen. Heat map indicates the number of retinal cells transduced (log10-transformed). The number of cells were averaged (excluding zeroes) across n=3 NHPs. Pan-retinal score indicates the number of retinal cell types transduced by a particular AAV variant. **d,** UMAP of AAV-transduced cells in fovea, periphery, and low-gate FACS-sorted retina (used to remove debris). **e,** Cryo-EM structure of ATX002, with the loop insertion highlighted in lime green.

Next, non-packaging variants were eliminated from the library: using a round of PCR, genomes from the packaged virus were amplified, recloned into the plasmid backbone, and re-packaged. The resulting subset of variants was named the ‘pro-pack’ library (fig. S1C). Both the TC6 and pro-pack libraries were injected intravitreally into non-human primate (NHP) eyes. One round of *in vivo* selection on TC6 and propack libraries was then performed to eliminate non-infectious variants.

To quantify AAV variant performance with single cell resolution, TC6, pro-pack, and *in vivo* selected libraries were screened through the scAAVengr-HUnT workflow. Relative representation of each variant in the library was determined through deep sequencing of the packaged variant pool prior to injection. The TC6 library was injected in 6 NHP eyes (n=6), the pro-pack AAV2-based library was injected into 9 NHP eyes (n=7), and the *in vivo* enriched AAV2-based library was injected into 3 NHP eyes (n=3) (Supplementary Table 1). Four weeks post-injection, retinas were isolated and dissociated into single cell suspensions, processed for single-cell RNA-seq, and sequenced. We analyzed 99,411 cells from TC6 AAV2-based library NHPs, 1,622,827 cells from pro-pack AAV2-based library NHPs, and 161,638 cells from *in vivo* AAV2-based library NHPs, as previously described in our low-throughput scAAVengr workflow^13^, with modifications to enable high-throughput library analysis. Briefly, retinal cell types were identified through cell type marker gene expression, and AAV transduction was quantified through GFP mRNA expression.

Single cell analysis identified 546 retinal-transducing variants from the TC6 library, 8,450 variants from the pro-pack library, and 1,178 variants from the *in vivo* library. Top candidate capsids were prioritized for further analysis based on packaging efficiency, changes in representation over library iterations, transduction across regions of the retina (central and peripheral), number of pan-retinal cells transduced and expression in specific cell types of interest (Fig. 1B, fig. S2).

An additional low-throughput scAAVengr run was performed using 233 variants that were selected for further analysis (Fig. 1C). AAV2, the parental serotype, and leading engineered vectors were also added as controls, including 7m8 and K912, which was developed in the context of canine eyes^13^. Variants were cloned and packaged individually with a ubiquitous CAG promoter driving an eGFP transgene containing a unique 25-bp barcode. Variants were titer-matched and pooled, then injected intravitreally into 6 NHP eyes (n=3) for a head-to-head *in vivo* analysis at higher resolution, using clinically relevant titers.

Analysis of the pooled AAV2-based variants at single cell resolution showed that the relative performance of AAV2, 7m8, and K912 was consistent with previous findings, with 7m8 and K912 outperforming the parental control. In addition, 72 novel scAAVengr-HUnT variants transduced greater numbers of cells than AAV2, and 27 variants outperformed 7m8 (Fig.1C, fig. S3A). In the peripheral retina alone, 124 variants outperformed 7m8 (fig. S3B). One AAV2-based variant in particular, called ATX002, containing amino acid insert sequence LAEHQTRPA, transduced the highest number of cells pan-retinally (Fig. 1D, Fig S3C), 14x more than 7m8, with consistent results across animals (Fig.1C,D). Of note, ATX002 efficiently transduced photoreceptors, indicating that it successfully crosses the ILM to reach the outer retina. In addition to penetrating retinal layers, ATX002 shows strong and consistent expression across central and peripheral retina.

### Capsid structure of ATX002

We compared the capsid of ATX002 with that of AAV2 (PDB: 8FYW) and 7m8 (PDB: 6U0R) to determine if the 9-residue insertion affected capsid assembly or structure. We generated a cryo-EM density map of ATX002 at 1.9Å resolution, from which we built an atomic model of the complete capsid (Fig.1E, EMDB: EMD-74043, PDB: 9ZCW). No electron density was observed for the loop inserts, indicating that they form flexible loops and do not interact with the rest of the capsid.

Comparison with the AAV2 and 7m8 structures indicates that the capsids are essentially identical (fig. S4, alpha-carbon RMSD of 0.772Å between VP3 proteins of ATX002 and AAV2, calculated using 515 shared residues), demonstrating that ATX002 functionality derives exclusively from the amino acid properties of the exposed insert sequence rather than structural changes.

### ATX002 outperforms current clinical-stage vectors

To validate pan-retinal performance, ATX002-CAG-mGL was injected intravitreally and bilaterally into n=4 NHPs. For comparison, 7m8-CAG-mGL was injected intravitreally and bilaterally into n=3 NHPs. Both vectors were injected at a dose of 1e11 vg/eye. ATX002 injections resulted in strong pan-retinal expression of mGL, across peripheral and central regions of the retina (Fig.2, fig. S5A). In contrast, 7m8 resulted in mGL expression that was markedly dimmer and limited to the foveal region of the retina. Cross-sectional imaging shows that ATX002 results in strong expression across all layers of the retina and transduces photoreceptors in periphery (Fig.2I,J) and fovea (Fig. 2K-M). A biodistribution study of ATX002 performed using multiplex ddPCR revealed that AAV genomes were restricted to the eye (fig. S5B).

**Fig. 2:**
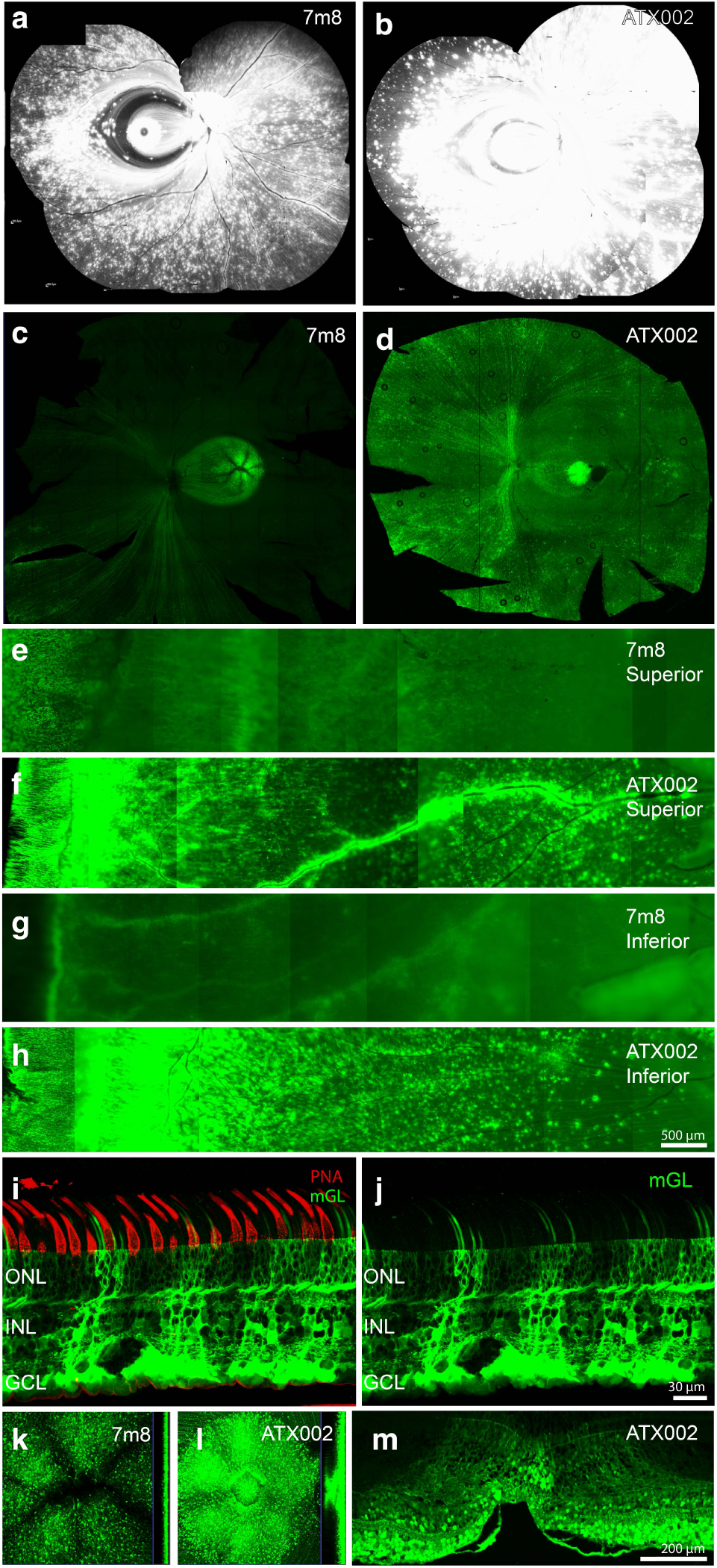
ATX002 expression in NHP retina following intravitreal injection. **a,b,** cSLO imaging of cynomolgus macaque eyes injected with 1E11 vg of 7m8 (A) or ATX002 (B). **c,d,** Retinal flatmounts from NHP eyes injected with 7m8 (C) or ATX002 (D). **e-h,** Montages of superior and inferior peripheral regions, from eyes injected with 7m8 or ATX002. **I,** Cross section of midperipheral retina from ATX002-injected eye, co-stained with PNA, a cone photoreceptor marker. **j,** Unstained cross section of midperipheral retina. **k,l,** Flatmounted foveas from 7m8 or ATX002-injected eyes. **m,** Cross section through fovea from an ATX002 injected eye.

We also tested the ability of ATX002 to deliver RS1, a therapeutic transgene, for the treatment of X-linked retinoschisis^14^, in NHP eyes. scRNA-seq showed expression in key retinal cell types, including photoreceptors and bipolar cells ^15^, following IVT injection. Compared to AAV8, recently used in IVT clinical trials for RS1^16^, ATX002 outperformed AAV8 by up to 328-fold, quantified by qRT-PCR (fig. S6B). These results suggest that ATX002 is an effective therapeutic delivery system at a significantly lower dose, potentially avoiding harmful immune response.

### ATX002 maintains retinal potency across species

We then tested ATX002 in mouse retina since preclinical studies are primarily carried out in rodent models. ATX002 (n=4), 7m8 (n=3), and AAV2 (n=3) were packaged with CAG-mGL and injected intravitreally into mouse eyes. Fundus imaging was performed, and retinal tissue was collected 4 weeks post-injection. Imaging revealed that ATX002 had markedly improved pan-retinal expression relative to both AAV2 and 7m8 (Fig.3A-F, fig. S6). These data indicate that ATX002 retains optimized fitness across species.

**Fig. 3:**
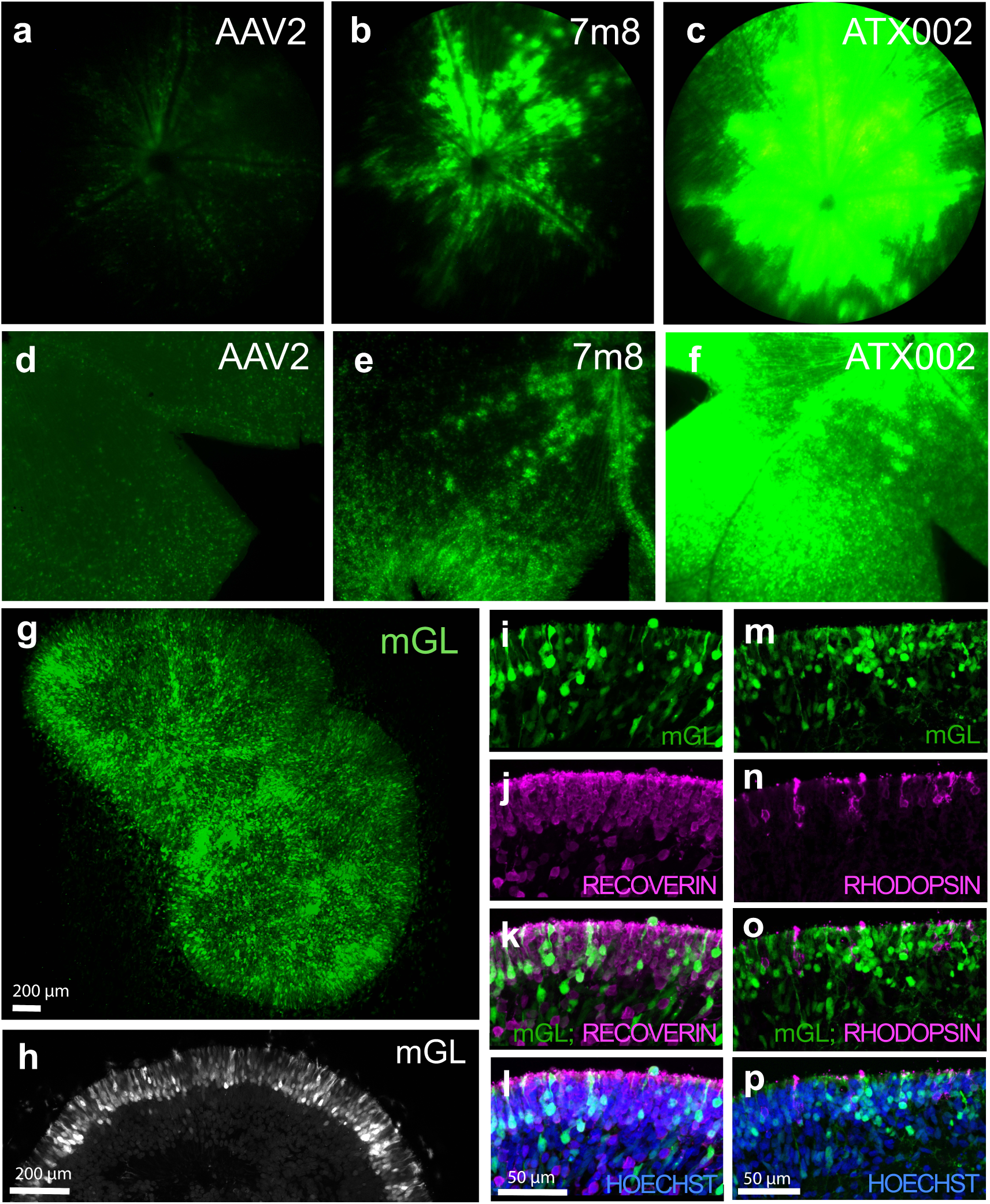
ATX002 expression in mouse retina and human retinal organoids. **a,b,c,** Fundus imaging in mice injected intravitreally with ∼3E10 vg of AAV2, 7m8 or ATX002. **d,e,f,** Flatmounts of retinas from injected mice. **g,** hiPSC-derived human retinal organoids transduced with ATX002-CAG-mGL, imaged as intact whole organoids by two-photon microscopy. **h,** Optical cross section of the organoids transduced with ATX002 depicted in G, highlighting the efficient transduction of the outer nuclear layer where photoreceptors reside. **i-p,** Confocal microscopy images from cross sections of an organoid transduced with ATX002, and co-stained with markers for photoreceptors (Recoverin, a generic marker, and Rhodopsin, a marker for rods) and nuclei (Hoechst).

To assess ATX002 performance in a human retinal model, we delivered ATX002 to iPSC-derived human retinal organoids (Fig. 3G-P, fig. S8). ATX002-CAG-mGL (1.8E11 vg in a 10 µL volume) was delivered in medium to human retinal organoids from late-retinogenesis to post-retinogenesis/”mature” developmental stages. Organoids were then fixed 21 days post-infection and imaged whole by two-photon microscopy (Fig. 3G,H) or cryosectioned and stained with antibodies for retinal cell types markers (fig. S7). Imaging revealed that ATX002 transduced photoreceptors efficiently, indicating that ATX002 tropism translates to human cells. ATX002 also outperforms 7m8 in iRPE culture, transducing up to 80% of iRPE cells 14 days post infection, dosing with 2E4 MOI (fig. S9).

### ATX002 outperforms AAV9 in mouse and NHP brain

We next tested whether the improved abilities of ATX002 extended to cells in the central nervous system (CNS). ATX002-EF1⍺-tdTomato and AAV9-EF1⍺-eGFP were co-injected via intracisternal magna (ICM) delivery into six male mice, aged 9-11 weeks. AAV9 was selected as a control as it is widely used for gene delivery to the brain. 10 µL of the AAV pool was injected at a rate of 1 µL/minute, delivering a total of 5e10 vg of each AAV. Tissues were collected four weeks post-injection and processed for histological analysis. ATX002 expression was higher than AAV9 in the cerebellum, thalamus, and brainstem (fig. S10A). In all brain regions imaged, ATX002 expression was largely neuronal, with few glial cells expressing tdTomato. In another group of mice, 5E10 vg of ATX002-EF1⍺-tdTomato and AAV2-EF1⍺-eGFP were co-injected via intraparenchymal injections to V1. Thirty days after injection, tissue was sectioned and imaged. ATX002 outperformed AAV2 in all mice tested (n=4) (fig. S10B).

We then evaluated ATX002 CNS transduction in NHP. ATX002-EF1⍺-tdTomato and AAV9-EF1⍺-eGFP were again titer-matched at 1E13 vg/mL and co-injected via ICM delivery into a male cynomolgus macaque between 2-3 years of age. A manual bolus of 1 mL of the final AAV pool was injected over the course of one minute, delivering a total of 5E12 vg of each virus. Tissues were collected four weeks post-injection, sectioned, cleared, and imaged with confocal microscopy.

Sagittal sections through the midline revealed markedly stronger expression mediated by ATX002 compared to AAV9 in regions across the brain including cerebellum, mammillary body, and anterior cingulate cortex (Fig.4A). Co-staining with a neuronal marker (Nissl stain) indicated that labeling was highly neuronal specific across brain regions (Fig.4B, fig. S11). Quantification of GFP or tdTomato labeled cells in sequential brain slices confirmed greater numbers of cells were transduced by ATX002 compared to AAV9 in all areas evaluated, including hippocampus, entorhinal cortex, substantia nigra, cerebellum, ventral tegmental area, hypothalamus, and anterior cingulate cortex (Fig.4C). Interestingly, ATX002 also labels large numbers of ependymal cells throughout the ventricles, central canal and cerebral aqueduct (fig. S11). Collectively, this data indicates that ATX002 possesses universally beneficial properties that improve performance across a range of species and tissues.

**Fig. 4:**
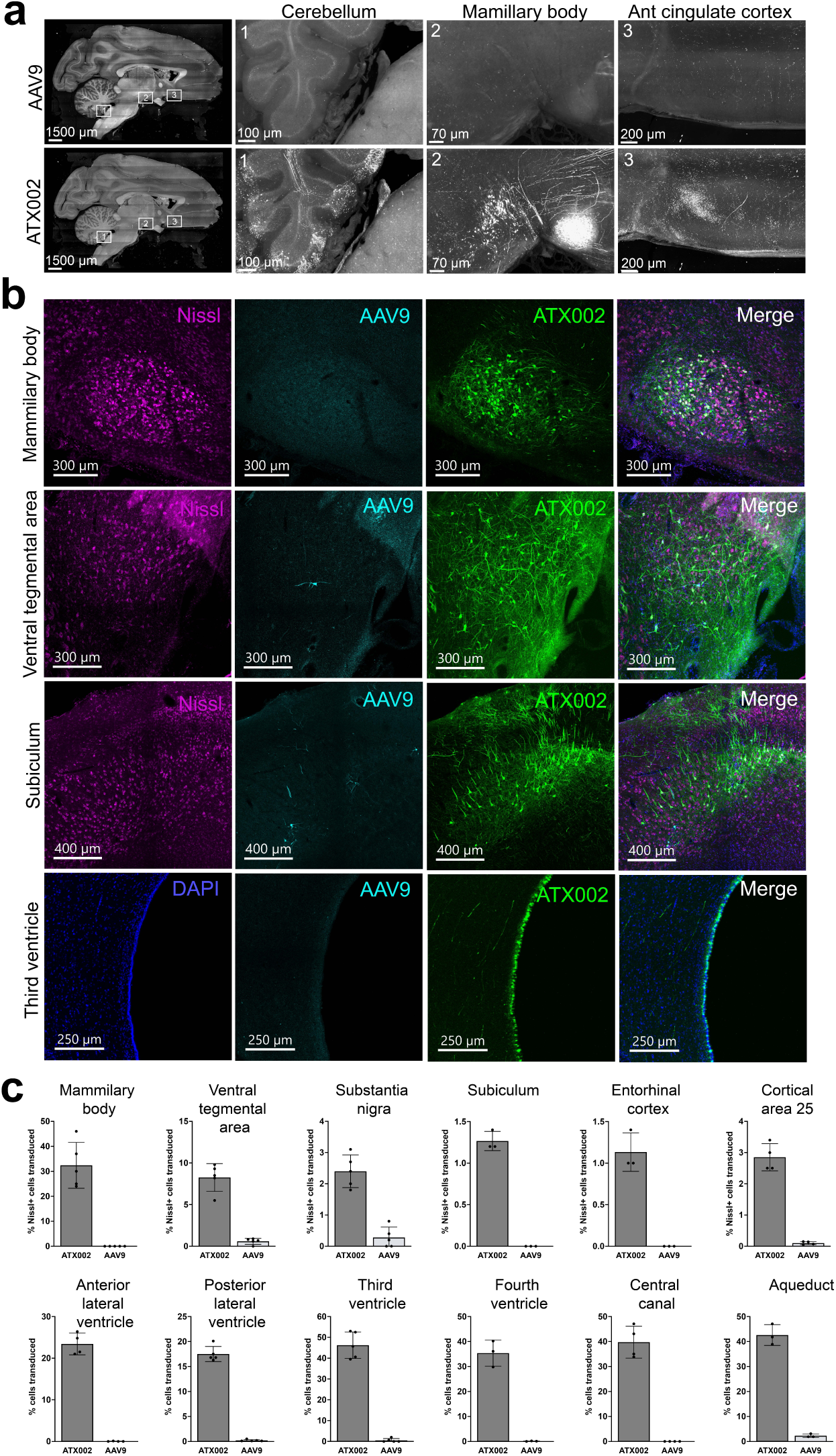
ATX002 expression in NHP brain following ICM injection. **a,** Imaging from a sagittal slice of NHP brain injected via ICM injection with ATX002-EF1⍺-tdTomato(shown in green) and AAV9- EF1⍺-GFP(shown in cyan). **b,** Co-labeling of NHP brain regions with DAPI (nuclei) and Nissl stain (neurons). **c,** Quantification of cell transduction across brain regions and in the ventricular system.

### Comparative structural dynamics analysis of AAV variants reveals bifunctional molecular mechanism underlying improved performance

Although considerable effort has been devoted to engineering novel AAV inserts to enhance transduction efficiency^17^, these efforts have largely lacked guidance from biophysical and mechanistic insights. Here, we present what is, to our knowledge, the first mechanistic investigation of engineered AAVs using MD simulations. We hypothesized that, for AAV2-based capsids, two major factors contribute to the fitness of a given AAV variant, including: (1) heparan sulfate binding affinity and (2) recognition and binding affinity to the AAV universal receptor (AAVR). To model these effects, we performed 1-µs all-atom MD simulations of AAV capsid segments consisting of 3 VP3 proteins at the 3-fold symmetry axis, both in complex with its receptor (to capture AAVR binding modalities) and in isolation (to model intraloop interactions in the absence of AAVR binding).

We selected five loop variants for MD simulations, in decreasing order of *in vivo* fitness: ATX002, 7m8, AAV2, var16 and ATX002_ins_589 (Fig. 5A). These variants were chosen to span a broad range of structural features and *in vivo* fitnesses as characterized by scAAVengr-HUnT: ATX002 as a new best-in-class vector, 7m8 as the current gold-standard, AAV2 as baseline, var16, which contains an insertion but performs equally to AAV2, and ATX002_ins_589, which underperforms relative to AAV2. Var16 has a relatively featureless insert sequence, composed mostly of nonpolar residues. ATX002_ins_589 is an insert-shifted version of ATX002, with the insertion located at position 589 instead of 588, and it was included to investigate the positional sensitivity of successful inserts.

**Fig. 5:**
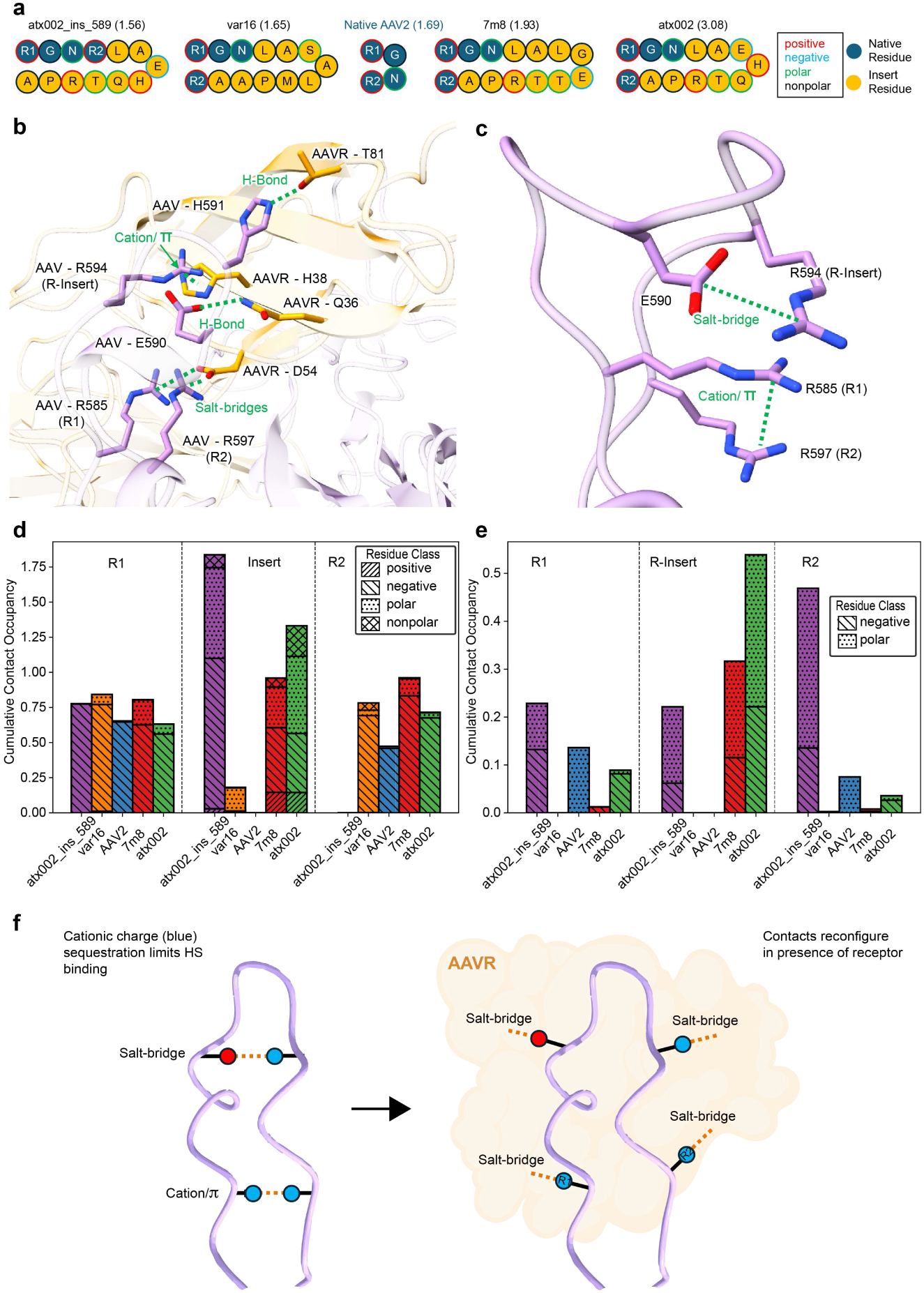
Comparative structural analysis of AAV variants. **a,** Loop insertion sequences for the variants included in the MD analysis, ordered by increasing *in vivo* fitness of retinal transduction as quantified by the scAAVengr-HUnT workflow (cells transduced, log transformed, in parentheses) **b,** Representative snapshot of an MD trajectory of the ATX002 (purple) – AAVR (orange) complex. **c,** Representative snapshot of a loop-alone ATX002 MD trajectory. **d,** Cumulative hydrogen bond contact frequencies between simulated variants and AAVR, binned as insert and native arginines. **e,** Cumulative hydrogen bond contact frequencies of insert and native arginines (when present) for isolated intraloop regions of simulated variants. **f,** Simplified schematic illustrating the bifunctional mechanism of ATX002. In the absence of AAVR (left), inserted charged residues form intraloop interactions that sequester charge and reduce nonproductive binding to heparan sulfate and other molecules. Upon AAVR engagement (right), these residues are repositioned to form favorable salt bridges and hydrogen bonds across the AAV–AAVR interface, enhancing receptor auinity and overall transduction euiciency.

MD simulations of AAV–AAVR complexes revealed key insights into the influence of loop insertions on AAV receptor engagement. First, simulations revealed that the two native arginines in AAV2, which we termed R1 and R2, form critical contacts with AAVR that are likely essential for functional binding (Fig. 5B,D, fig. S12-S16). Interactions with AAVR occur through salt bridges between the arginines in AAV2 and a glutamic acid on AAVR (at position 54 in the reference system, Fig. 5B). For engineered variants that outperform AAV2, these interactions with AAVR are increased in frequency in our MD simulations. While both 7m8 and ATX002 (Fig. 5D) show similar interactions between R1 and AAVR compared to AAV2, interactions between R2 and AAVR increased, likely contributing to their increased *in vivo* fitness. In contrast, for insert-shifted ATX_ins_589, these interactions are disrupted, and the displaced arginine R2 can no longer form a stable contact with D54 or any other residue in the AAVR, likely resulting in much weaker affinity to the receptor, and explaining the poor *in vivo* performance of this variant.

In addition to R1 and R2 contacts, the introduction of additional loop residues allows the capsid to extend its interaction interface, improving surface complementarity and enabling more effective engagement with AAVR. Specifically, the introduction of charged residues in 7m8 and ATX002 presents an advantage for AAVR binding. Both 7m8 and ATX002 contain an inserted arginine and glutamate that form productive interactions with AAVR, likely contributing to enhanced receptor binding (Fig.5B). MD trajectories show the inserted arginine forming salt bridges with both a glutamate and an aspartate on the AAVR. The incorporation of other positively or negatively charged and polar residues in the loop insertions further enhances hydrogen bonding and electrostatic interactions across the interface (Fig.5D). These expanded networks appear to stabilize the complex and contribute to the improved receptor affinity observed in variants that outperform AAV2. In contrast, var16, composed primarily of nonpolar residues that cannot form these interactions, shows neither enhancement nor impairment of *in vivo* fitness relative to AAV2.

While the charged and polar amino acids in the inserts of these high-performing AAV variants are favorable to AAVR binding, we also investigated their effect in earlier steps of the AAV transduction pathway. In particular, we wanted to understand the potential effect of the inserted arginine, which is present in both ATX002 and 7m8, on HS binding. Reduced HS affinity is thought to be a desirable property for AAV2 vectors in ocular gene therapy^18^. Strong HS binding could result in the increased unproductive retention of AAVs in barriers such as the HS-rich ILM, preventing transduction of targeted cells. HS is known to bind to arginines R1 and R2 at the base of the VR-VIII loop^19^, and the proximity of the additional positively charged arginine in these engineered variants could increase this affinity. Surprisingly, despite containing this additional arginine, 7m8 has been shown to have lower affinity for HS than AAV2^12^. To understand these mechanisms, we performed 1-µs MD simulations on the same 3xVP3 segments of all five variants, in the absence of AAVR.

For ATX002, simulations revealed that the inserted arginine strongly interacts with neighboring polar and negatively charged residues through salt bridges with the inserted glutamate (Fig.5C,E, fig. S17-S20). This charge pairing effectively shields these residues from unwanted interactions with molecules in the ocular milieu, such as the interaction between the inserted arginine and HS. Simulations of 7m8 show a similar mechanism though of lower magnitude than ATX002 (Fig.5E). The endogenous R1 and R2 arginines in both variants remain available for HS binding, though this interaction may be reduced relative to AAV2 through steric occlusion from the inserted residues (Fig.5E).

In ATX002_ins_589, the inserted arginine is shielded by the inserted glutamate, similar to 7m8. However, its endogenous arginines, and R2 in particular, are strongly involved in interactions with negative and polar residues. While limited HS affinity is a desired property, too low of an affinity would likely have a deleterious effect as it would result in insufficient binding to the surface of targeted cells. Furthermore, this could also contribute to the lack of interactions between the displaced R2 and AAVR.

## Discussion

Here, we describe a complete platform for the engineering of new, potent gene therapy vectors with high clinical potential. We used this workflow to develop best-in-class AAV vectors for retinal gene therapy, validate these variants across species and tissues, and to investigate the mechanism underlying improved capsid fitness using structural dynamics.

The scAAVengr-HUnT workflow provides AAV capsid screening of large libraries with single-cell resolution on a high-throughput scale, and identifies high performing engineered AAV variants on the basis of mRNA expression. Importantly, this pipeline allows for efficient selection performed *in vivo* in NHPs, which are a critical large animal model for the retina. NHP retinas are anatomically and structurally similar to humans, containing a fovea, a thicker inner limiting membrane than in rodents, and substantial vitreous humor. Optimization and quantitative analysis in the context of NHP retina provides confidence that these AAV variants will perform similarly in humans. However, the scAAVengr-HUnT workflow may be adapted to any species, tissue, or cell type for which marker genes are available.

The top-performing AAV variant, ATX002, was identified as a top pan-retinal performer across both the fovea and periphery, outperforming clinical-stage 7m8 by more than 1log10 and parental AAV2 by 2log10. ATX002 is therefore a strong candidate for treating retinal diseases that require widespread distribution of therapeutic transgene, or delivery to the outer retina. Data from RS1 delivery experiments demonstrate up to 300-fold improvement in therapeutic transgene expression compared to wildtype AAV8, currently in clinical use. Importantly, ATX002 retains its potency across mice, NHPs, and human organoid retina, indicating that this capsid will provide reliable performance across preclinical animal models and human patients. Additionally, ATX002 shows promise for applications in neuronal tissue outside of the retina, transducing brain from ICM and IPa injection. Together, these results highlight the universality of capsid fitness and suggested to us a mechanism involving ATX002’s main receptors. We therefore turned to a biophysical approach to investigate the mechanism underlying ATX002 infectivity.

We used MD to better understand the impact of different amino acid inserts on capsid performance. These molecular simulations offer the first mechanistic study at the atomic level of engineered AAV variants and suggest a novel and general bifunctional mechanism: the inserts of high performing variants incorporate charged and polar residues that lead to additional favorable interactions with the AAV receptor but are otherwise shielded to prevent AAV retention through unwanted interactions with HS or other barriers (Fig. 5F). The degree to which these mechanisms are successfully exploited correlates with the range of *in vivo* fitness demonstrated by these different AAV variants. These findings underline the multistep pathway that AAV must traverse to successfully deliver a therapeutic transgene and highlight that capsids must possess multiple abilities to achieve optimal fitness, including efficient receptor binding and extracellular trafficking. Importantly, this work establishes novel approaches for future AAV engineering and highlights the limitations of methods based on sequence space alone. Molecular dynamics is a previously untapped approach in the computational design of improved AAV vectors. We predict that expanding current sequence-based ML methods with molecular dynamics features will unlock powerful new strategies for the design of safe and efficient next-generation gene therapy vectors.

## Acknowledgements

We thank Jackie Breter for assistance with sample collection. We thank Stacy Cashman and Nicole McLane, as well the University of Pittsburgh DLAR staff for assistance with animal surgeries. We thank Fabian Rueckert, Olivier Partouche, and Erich Kueng for technical assistance. We thank Jane Opgaard, Lora Waybright, and Simon Bassett for project management. We thank Yong Zheng for providing the RS1-myc tag construct design. Figures were made with Biorender (https://BioRender.com).

## Author contributions

MB and LCB conceived the project. MEJ, ML, and MB performed computational analysis of single-cell RNA-Sequencing. BEO, HNJ, MS, LC, JEH, BH, MG, and LCB performed molecular biology experiments including scRNA-Seq and imaging. HNJ and JEH carried out CNS studies. THT and MB performed structural dynamics analysis. JFC performed cryo-EM experiments and JFC and MB analyzed the data. HA, WRS and LCB performed surgical procedures. JK, IC, SS, FR, RS, FK, SF and JK performed retinal validation experiments on ATX002. KF, SF, and SdS performed experiments in human retinal organoids. PAS, JAS, and RL provided funding, support, and supervision. MEJ, BEO, MB and LCB wrote the manuscript with contributions, review and editing from all authors.

## Data and materials availability

The ATX002 capsid structure and sequence is available through PDB: 9ZCW. Data from figures is available on Dryad. All algorithms and tools required to reproduce findings are described in the main text or “Methods”.

## List of Supplementary Materials

Materials and Methods Figs S1-S20

Tables S1 and S2

## Methods

### Study approval

All intraocular procedures performed on mice were in accordance with the ARVO statement for the Use of Animals in Ophthalmic and Vision Research and approved by the University of Pittsburgh IACUC committee (IACUC #21058910). Intraocular procedures conducted on primates were in accordance with the ARVO Statement for the Use of Animals and the guidelines and were performed with approval from the Division of Laboratory Animal Resources at the University of Pittsburgh (IACUC #21058910). ICM and IPa injections in NHPs and mice were conducted in accordance with the ARVO Statement for the Use of Animals and the guidelines and were performed at Northern Biomedical Research. For NHP experiments performed at NBR, the Animal Care and Use Protocol (179-002A) was approved by the Institution Animal Care and Use Committee in compliance with the Guide for the Care and Use of Laboratory Animals, DHHS (NIH), No. 86-23 and the Animal Welfare Act (9 CFR 3).

### Data analysis

All measurements were taken from distinct samples.

### Primates

Cynomolgus macaques and rhesus macaques were used for all NHP studies. Animals were between 2–10 years of age, and IVT injections were performed using previously described methods^13^. All NHPs used in these studies were screened for neutralizing antibodies and had titers of ≤1:10. Monkeys received daily oral doses of cyclosporine (6 mg/kg) and prednisone (1 mg/kg) for the duration of the study. At the conclusion of the experiment, euthanasia was carried out with an IV overdose of sodium pentobarbital (75 mg kg−1), as recommended by the Panel on Euthanasia of the American Veterinary Medical Association. For single-cell RNA-Seq experiments, animals were perfused with ACSF prior to enucleation. A summary of minor adverse events related to the procedures is summarized in Supplementary Table 1. Other than uveitis in some eyes, no other adverse events were noted.

### AAV packaging

AAV vectors were produced in HEK293T cells (ATCC), or 293AAV cells (Cell Biolabs) using a double or triple transfection method ^21^. Short tandem repeat profiling was done by ATTC Cell Line Authentication Service and all cell lines were checked for mycoplasma using Hoechst staining.

Recombinant AAVs were purified by either iodixanol gradient ultracentrifugation or affinity chromatography on an FPLC using an AVB column (Cytiva) and then buffer exchanged and concentrated with Amicon Ultra-15 Centrifugal Filter Units (#UFC8100) in DPBS and titered by quantitative PCR relative to a standard curve using ITR-binding primers or by using QuickTiter AAV Quantitation Kit (Cell Biolabs).

### Construction and characterization of AAV libraries

AAV libraries were constructed by cloning the P40 promoter upstream of VP1 (fig. S1A). A 6-amino acid insertion, surrounded by a leucine-alanine linker upstream and an alanine downstream, was placed at position 588 of VP1. Libraries were constructed using trimer synthesis, such that all amino acids were represented at all 6 amino acid positions. Following the stop codon of VP1, a ubiquitous CMV promoter drove expression of an eGFP gene, which was following by a random 20-bp barcode and a bgh-PolyA. Barcodes were generated with random nucleotides at all 20 positions. In order to determine the GFP barcode-AAV capsid variant pairings, the region of the plasmid including the capsid variant and GFP barcode was PCR amplified, ligated, and deep sequenced; These are referred to as dictionaries in the scAAVengr pipeline. The representation of variants in the library was also determined, by PCR amplification and deep sequencing of 20-bp barcodes taken from the packaged virus of the injection pool.

### Propack library construction

To remove non-packaging vectors from the library, one round of library packaging was performed, followed by PCR amplification of VP1, GFP and 20-bp barcodes. These amplicons were re-cloned into the AAV vector backbone, and re-packaged for injection into primate eyes. The balance of variants in this library was determined through PCR amplification and deep sequencing of 20-bp barcodes.

#### *In vivo* enrichment

To enrich for variants in the library with the ability to transduce retinal cells following IVT injection, PCR amplification of VP1, GFP and 20-bp barcodes was performed following injection of the libraries into primate eyes. Retinal and RPE samples from regions across primate eyes were amplified, and these amplicons were re-cloned into the AAV vector backbone and re-packaged for injection into primate eyes. Following cloning, libraries resulting from retinal regions were pooled evenly. The balance of variants in this library was determined through PCR amplification and deep sequencing of 20-bp barcodes.

### Low-throughput library construction

AAV2 capsid variants identified as top performers in scAAVengr-HUnT were subsequently packaged and purified for a low-throughput scAAVengr analysis. Unique 8-bp DNA barcodes were cloned after the stop codon of eGFP, in an AAV ITR-containing plasmid construct containing a self-complementary CAG promoter driving eGFP expression (scCAG-eGFP-Barcode-bghPolyA). *Cap* genes were cloned into a separate AAV rep/cap plasmid (Addgene #64839). Individual AAV variants were then packaged separately with constructs containing different barcodes using a triple transfection method in HEK293 suspension cells. Following packaging, variants were pooled together for purification on an AKTA Pure 25M (Cytiva, Marlborough, MA) using a Repligen AAV2 affinity column (Repligen, Waltham, MA). After purification, pools were sequenced to ensure that the relative abundance of each vector in the pooled library was equal. Each variant was amplified using primer/adapters and sequencing on an iSeq flow cell. Once the relative abundances of each variant were confirmed to be equal, pools were titered using the qPCR AAV Titer Kit (ABM, Bellingham, WA). Titers for all variants in the pool were determined to be within ±1 log from the average variant in the pool, a range that was found to be compatible with accurate normalization across samples.

### Single-cell dissociation of primate retina

The NHP retinas were dissected, and regions of interest were isolated (macula, superior, and inferior periphery). Retinal tissue was placed in Hibernate solution (Hibernate A -Ca Solution, BrainBits LLC), and cells were then dissociated using Macs Miltenyi Biotec Neural Tissue Dissociation Kit for postnatal neurons (130-094-802) according to manufacturer’s recommendations. Dissected retina pieces were incubated with agitation at 37 °C and further mechanically dissociated. The dissociated neural retina was filtered using a 70 μm MACS Smart Strainer (Miltenyi Biotec) to ensure single-cell suspension. Cells were resuspended in 0.1% BSA in D-PBS and processed immediately for scRNA-seq.

### FACS

Following dissociation, a Miltenyi MACS Tyto sorter was used to remove dead cells and debris. Cells were resuspended in 0.1% BSA in D-PBS and processed immediately for scRNA-seq.

### Single-cell RNA-seq of primate retina

Rhesus macaque and cynomolgus macaque samples were prepared for single-cell analysis using a 10x Chromium Single Cell 3’ v3.1 kit. Briefly, single cells from retina samples were captured using a 10x Chromium system (10x Genomics), the cells were partitioned into gel beads-in-emulsion (GEMS), mRNAs were reverse transcribed and cDNAs with 10x Genomics Barcodes were created with unique molecular identifiers (UMIs) for different transcripts. Purified cDNA was PCR amplified and further purified with SPRIselect reagent (Beckman Coulter, B23318). Final libraries were generated after fragmentation, end repair, A-tailing, adaptor ligation, and sample index PCR steps according to 10x Single Cell 3’ workflow. Targeted gene enrichment was run on these 10x-prepped cDNA samples using 10x Target Hybridization Kit (PN-1000248) and 10x Library Amplification Kit (PN-1000249). IDT xGen Custom Hybridization Panel containing biotinylated probes for retinal cell type marker genes and GFP was used to pull down cDNA of interest. 10x libraries were pooled and all samples were submitted for deep sequencing on Illumina Novaseq S4 flowcells at the UPMC Genome Center. Sequencing depth was targeted at 20,000 reads per cell for the standard scRNA-seq analysis and between 5,000 and 10,000 reads per cell for the targeted gene enrichment analysis.

### Single-cell RNA-seq pre-processing

Sequencing data was demultiplexed into sample-level fastq files using Cell Ranger mkfastq (v3 10x Genomics). For full transcriptome samples, alignment and cell demultiplexing were run using STARsolo [17] (v2.7) with default parameters. DropletUtils [18] (v1.4.3) was used after STARsolo to remove empty droplets (lower.prop = 0.05). Cynomolgus macaque samples were aligned to the Macaca_fascicularis_5.0/macFas5 reference obtained from UCSC. Gene annotation for the cynomolgus macaque was created by lifting over the pre-mRNA gene annotations from the hg38 Ensembl human genome. Doublets (10x droplets containing two cells instead of one) were then identified using SCDS [19] (v1.0.0). Any droplets with a hybrid score >1.3 were considered doublets. CellRanger (v6.0.1) was run to align and demultiplex cells using the targeted gene panel and human reference GRCh38. Reads were normalized and log-transformed in Scanpy [20] (v1.4.4.post1).

### Single-cell RNA-seq cell type identification

Scanpy (v1.4.4.post1) was used for the analysis of both full transcriptome and target gene enriched scRNA-seq samples. Principal component analysis (PCA) was used to embed into low dimensional space. Harmony was used to integrate samples across NHPs and 10x processing methods (full transcriptome and target gene enrichment)^22^. Normalized counts were saved as raw data and used for differential gene expression analysis. Leiden clustering was run to identify cell clusters. Cell types were determined by running a differential gene expression analysis using Scanpy’s ‘rank_gene_groups’ function. We used a hypergeometric test and calculated the significance of the intersection of marker genes from one cluster with the published marker genes of each retinal cell type. A Bonferroni p-value correction was applied to account for multiple-hypothesis test. Each cluster was assigned a cell type based on the most significant marker gene intersection p-value.

For clusters where the hypergeometric test could not identify a specific cell type match, we annotated the cell type based on marker gene expression using a known cell type marker database23-25.

### Single-cell AAV transgene and capsid quantification

The performance of AAV capsid variants was analyzed based on quantification of AAV variant-mediated GFP-barcode mRNA expression. GFP barcodes were quantified using Salmon (v1.4.0) with a custom GFP library-based reference. Quantification was further refined using Seqkit grep to search for appropriate surrounding sequence. GFP reads were mapped to their appropriate 10x cell barcodes using paired-end sequencing information. GFP counts were further refined by collapsing and summing the unique molecules (UMIs) within a cell. A cell x gfp gene matrix was then created and mapped to the identified cell types using the 10x cell barcodes. For high-throughput libraries, GFP barcodes were mapped to their respective capsid variants using a ‘dictionary’ dataset – a separate deep-sequenced run of library plasmid pools containing the GFP barcodes and AAV capsid variant pairings. Representation of each capsid variant was quantified using the GFP barcodes from a deep-sequenced run of packaged AAV taken from the injection pool. For low-throughput library, GFP barcode and AAV capsid variant pairings were known a priori and each GFP barcode was mapped to its assigned capsid variant.

### Electron microscopy and structure determination

Particle concentration and assembly state were assessed by negative stain electron microscopy. 2.5µL of purified sample was pipetted on to a freshly glow discharged copper grid with a continuous film of graphitized carbon, washed in 1% uranyl acetate negative stain, and imaged in a TFS Tecnai TF20 transmission electron microscope (Thermo Fisher Scientific, MA, USA) operating at 200 kV and equipped with a TVIPS XF416 CMOS camera (TVIPS GmbH, Gilching, Germany). Images were collected with the TVIPS *EMplified* software.

A data set for structural analysis was collected on a TFS Krios 3Gi cryo-electron microscope (cryo-EM) equipped with a TFS Selectris energy filter and a Falcon 4i direct electron detecting camera. 2.5µL of purified sample was pipetted on to a freshly glow discharged Quantifoil R2/1 grid (Quantifoil Micro Tools GmbH, Großlöbicha, Germany) and then blotted and plunge-frozen into a liquified ethane:propane mix at a 60:40 ratio^26^ using a TFS Vitrobot Mk 4. Grids were mounted in the Krios and imaged at 300 kV under control of the TFS *EPU* v3.7 software at a magnification of 165,000x, corresponding to 0.72 Ångstroms/pixel at the sample, and a total dose of 30 e/Å^2^. 10377 movies were collected in electron counting mode and with an energy filter slit of width 10 eV. Aberration-free image shift (AFIS) was used to improve the data collection rate.

The cryo-EM dataset was analyzed with RELION 5.0^27^, including motion correction, contrast transfer estimation and refinement according to AFIS groups, classification, orientation refinement and particle polishing. Due to modest sample concentration, 14969 images of full capsids and 14802 images of empty capsids were selected. Exhaustive refinement with icosahedral symmetry imposed yielded a density map at 1.89 Å resolution and 1.94 Å resolution, respectively, at the commonly accepted Fourier shell correlation limit of 0.143, according to the “gold-standard approach” implemented in RELION ^28^. Ewald sphere correction^29^ was used to improve the resolution estimate by 0.05 Å in each case. An initial capsid model was developed with ModelAngelo^30^ and further refined in Phenix 1.21^30^.

### Fluorescent immunohistochemistry and imaging of mGreenLantern in NHP eyes

Eyes were enucleated and limbus was marked at the 9 o’clock and 12 o’clock positions using ink. Eyes were fixed in 10% neutral buffered formalin at ambient condition for 24 hours for eye cross-section IHC. For flat-mount IHC, eyes were fixed in 4% PFA (paraformaldehyde) at 4°C for 8 to 10 hours. Formalin-fixed paraffin embedded eyes were cross-sectioned to 5μm thin sequential sections mounted on glass slides. Samples were immunostained using Ventana protocol including deparaffinization, antigen retrieval in citrate buffer, blocking, 1h incubation with primary antibodies at 37°C, and 1h incubation with secondary antibodies at room temperature following manufacturer’s instructions. Reporter mGreenLantern transgene was stained with primary anti-GFP (Invitrogen) and host-compatible secondary antibodies. Nuclei were stained with DAPI. Control staining was performed without primary antibody. Images were captured using Olympus VA120 Slide Scanner with 10x objective. PFA-fixed eyes were dissected with preserved orientation marks to obtain central, nasal, temporal, superior, and inferior quadrants. Images of flat-mounted retina were acquired using high resolution confocal Zeiss LSM 710 microscope (Carl Zeiss Microscopy GmbH) with 10x or 20x objective and tiled z-stack scan.

### qRT-PCR in NHP tissue

A qRT-PCR assay was performed to assess the expression fold change of the RS1 transgene delivered with ATX002 or AAV8 capsids. Tissue samples from the nasal inferior and temporal superior regions of NHP left and right eye retinas were collected. The tissues were sectioned into far periphery, mid periphery and center, then frozen at -80C. Total RNA from tissue samples was extracted following the AllPrep DNA/RNA kit from Qiagen (cat. 80204, Germantown, MD). The Verso Reverse Transcriptase protocol (cat. AB1453A, Thermo Fisher Scientific, Waltham, MA) with anchored oligo-dT primers, was used to generate cDNA from the RNA extracts. Lastly, q-PCR was performed following the PowerUp SYBR Green Master Mix for real-time PCR protocol from Applied Biosystems (cat. A25742, Waltham, MA) on the QuantStudio3 by Applied Biosystems (Waltham, MA). Relative expression of RS1 transgene was calculated using the double ΔCt method, with normalization to the housekeeping gene *GAPDH*.

### iRPE culture

Human iPS-derived human RPE cells (iCell Retinal Pigment Epithelial Cells Lot 106881, FujiFilm CDI) were expanded to confluence on laminin-521-coated 96-well plates using RtEGM Retinal Pigment Epithelial Cell Growth Medium Bullet Kit (Lonza) according to the manufacturer’s instructions. Differentiation was induced by serum-depriving the confluent cultures for a period of 2-4 weeks. AAV transduction efficiency was determined by in-life imaging (Opera Phenix) using digital phase contrast and a segmentation protocol.

### Biodistribution studies following intravitreal injections

Genomic DNA was isolated from the collected tissue samples using a commercially available DNA extraction kit (n=3). The quantity and purity of the extracted DNA were measured using a Nanodrop spectrophotometer and Agilent TapeStation instrument. Viral DNA biodistribution was quantified by multiplex ddPCR using 10 ng of input total DNA. The housekeeping gene RPP30 served as normalization control.

### Human induced pluripotent stem cells (hiPSC) maintenance

The control hiPSC line IMR90-4 (WiCell) was utilized for differentiation of human retinal organoids. Following WiCell instructions, hiPSCs were cultured at 37°C and 5% CO2 in a humidified incubator using mTeSRTM plus medium (STEMCELL Technologies, #100-0276) in 6-well plates (Corning, 3516) coated with hESC-qualified Matrigel (Corning, 354277). hiPSCc were passaged at 80% confluency, about every 4-5 days.

### Differentiation of human retinal organoids

The organoid differentiation procedure was based on previously described protocols^31,32^. Briefly, hiPSC were dissociated to single cells using Accutase (Thermofisher Scientific, # 00-4555-56) and mechanical dissociation with a P1000 in 1ml of mTeSRTM plus containing 10uM Rock inhibitor Y-27632 (STEMCELL Technologies, #72304). hiPSC were then plated on 100mm petri dishes (VWR # 25384-088) with a total volume of 10ml of mTeSRTM plus to promote formation of embryoid bodies (EBs). On Day (D) 1, approximately 1/3 of the medium was exchanged for neural induction medium (NIM) containing DMEM/F12 (Gibco, #11330057), 1% N2 supplement (Gibco, #17502048), 1x NEAAs (Sigma, #M7145), and 2mg/ml heparin (Sigma, # H3149). On D2, approximately 1/2 of the medium was exchanged for NIM. On D3 EBs were plated in new 60mm petri dishes (Corning #430166) in NIM. Half of the medium was changed every day until D6. At D6, BMP4 (R&D, #314-BP) was added to the culture at a final concentration of 1.5nM. At D7, EBs were transferred to 60mm dishes previously coated with Growth Factor-Reduced (GFR) Matrigel (Corning, #356230) and maintained with daily NIM changes until D15. On D16, NIM was exchanged for retinal differentiation medium (RDM) containing DMEM (Gibco, #15140122) and DMEM/F12 (1:1), 2% B27 supplement (without vitamin A, Gibco#12587-010), NEAAs, and 1% penicillin/streptomycin (Gibco, #15140122). Feeding was done daily until D27. On D28, EBs were dislodged from the plate by checkerboard scraping, using a 10µl or 200µl pipette tip. Aggregates were washed three times and then maintained in 6 well culture plate (Greiner bio-one, #657185) in RDM medium. The medium was changed every 2-3 days until D41. During this time, organoids were sorted based on their morphology under a stereoscope with transmitted light. Organoids were clearly identified based on the presence of a typical phase-bright, pseudo-stratified neuroblastic epithelium. On D42, the medium was changed to RDM plus, which consisted of RDM supplemented with 10% FBS (Gibco, #16140071), 100µM Taurine (Sigma, #T0625) and 2mM GlutaMax (Gibco, #35050061). Feeding was done every 2-3 days, depending on the color of the medium. At D92, B27 supplement in RDM was switched to N2 supplement (Gibco, #17502-048).

### Organoid infection with AAVs

Organoids at D98 (W14), D114 (W17) and D294 (W42), covering a range of stages from late neurogenesis to “mature” stages and derived from different initiation batches, were infected with ATX002 (total n = 9 organoids). Briefly, a single organoid was transferred to one well of a 48-well plate with 600µl of medium and 10µl of ATX002-CAG-mGL, 1.8E13 vg/mL, was added to the medium after a quick spin, either combined or individually. The medium was changed regularly based on its color, typically once per week. Organoids were then fixed 21 days after infection and further processed for immunohistochemistry in cryo-sections or imaged as intact whole organoids by two-photon microscopy.

### Immunostaining of organoid cryo-sections

Human retinal organoids were fixed with 4% PFA at RT for 30 min, washed 3 times with PBS, and cryoprotected in 1x PBS-30% sucrose at 4°C O/N. The next day, organoids were pre-embedded in a mixture of OCT and PBS-30% sucrose (1:1) for 30 minutes to 1 hour on ice and embedded in individual blocks in the same mixture in dry-iced 90% ethanol. Cryosections of 16µm thickness were alternated across Superfrost Plus slides (Fisher, #12-550-15). Once fully dried, slides were stored at -80°C until further processing. On the day of staining, slides were removed from -80°C and allowed to warm up to RT for approximately 20min. Slides were washed 3 times with PBS, and blocked 1h at RT with 5% Donkey Serum and 0.1% Triton X-100 in 1x PBS. Primary antibodies were diluted in 2% Donkey Serum in PBS and incubated at 4°C O/N. Primary antibodies used included anti-Pax6 (#BDB561462, 1:1000 dilution); anti-OTX2 (R&D, #AF1979SP, dilution 1:50); anti-RECOVERIN (Chemicon, #AB5585, dilution 1:500); and anti-RHODOPSIN (Millipore, cat# MAB5356, 1:100 dilution). Slides were rinsed 3 times, 5min each, with 1x PBS and incubated with appropriate donkey raised Alexa-Fluo secondary antibodies for 1 hour at RT and counterstained with Hoechst (Thermofisher, catalog #62249. Slides were then mounted with Fluoromount-G solution (Southern Biotech, #0100-01) and imaged using a Olympus fv 1200 Confocal with 40x, NA1.3 objective.

### Whole organoid imaging by two-photon microscopy

Mesoscopic fluorescence imaging of CUBIC-based tissue cleared GFP-expressing organoid was performed using a point-scanning two-photon excitation fluorescence microscope (Ultima 2P Plus, Bruker, WI). Images were acquired at a resolution of 2048 × 2048 pixels through a 16× water-immersion objective (Nikon 16×/0.8 NA) with zoom at 0.8x (overscan mode) achieving a 1.42x1.42mm field of view (0.68 µm/pixel). A full volumetric dataset of each organoid was obtained by collecting sequential optical sections over a total imaging depth of 750 µm with 1 µm steps in the axial (z) dimension. Two-photon excitation was provided by a tunable femtosecond pulsed infrared laser (Insight X3, Spectra-Physics, CA), set to 965 nm for optimal excitation of GFP. The laser beam was directed into the microscope via a dichroic mirror (ZT1040dcrb-UF3, Chroma, VT), and scanning was performed in galvo–galvo mode. The microscope was controlled using PrairieView software control (vX5.5, Bruker, WI), with a dwell time of 6.2 µs/pixel and a photomultiplier tube (PMT) gain setting of 600 HV. All z-stacks were acquired sequentially and subsequently processed for three-dimensional visualization and quantitative analysis in ImageJ.

### Intraocular injections in mice

Adult wild-type C57Bl6J mice (Jackson Labs) were used for all injections. Before vector administration, mice were anesthetized with 100 mg/kg Ketamine, 20mg/kg Xylazine by intraperitoneal injection. An ultrafine 30 1/2-gauge disposable needle was passed through the sclera, at the equator and next to the limbus, into the vitreous cavity. Injections were made with direct observation of the needle in the center of the vitreous cavity. The total volume delivered was 1 µl.

### Mouse Intracisterna Magna (ICM) injection

Prior to the procedure, 1 mg/kg slow-release buprenorphine was subcutaneously administered to the animal to provide 72h of analgesia. For the ICM injection, animals were deeply anesthetized with 2.5% isoflurane and placed in the prone position in a stereotaxic head holder, with the head and body at an angle of 120°. The skin and muscle at the nape were retracted and a 36G Hamilton syringe was used to pierce the dura mater at the site of the cisterna magna. 10 µL of an AAV pool (ATX002+wtAAV9) was injected at a rate of 1 μL/min. Following the injection, the needle remained in position for ten minutes to avoid the potential for backflow. The needle was then retracted, skin was sutured, and animals were allowed to recover on a warming mat.

### Mouse Intraparenchymal (IPa) injection

Prior to the procedure, 1 mg/kg slow-release buprenorphine was subcutaneously administered to the animal to provide 72h of analgesia. For the IPa injection, animals were deeply anesthetized with 2.5% isoflurane. The scalp was shaved and prepped using 0.5% chlorhexidine and 70% ethyl alcohol, and the animal was placed in a stereotaxic head holder (David Kopf Instruments, Tujunga CA). The skin was retracted and a high-powered drill was used to perform a craniectomy to expose the cortical surface of the left hemisphere. A Hamilton syringe was lowered into V1 (coordinates: Posterior: 2.8mm; Left: 2.4mm; Deep below the skull: 1mm). Using a microinjection unit (David Kopf Instruments), 1 µL of an AAV pool (ATX002+wtAAV9) was injected at a rate of 0.15 µL/min.

Following the injection, the needle remained in position for ten minutes to avoid the potential for backflow. The needle was then retracted and the craniectomy covered with bone wax (Ethicon). A local anesthetics mixture of bupivacaine and lidocaine applied to the skin at the site of the craniectomy prior to suturing overlying skin. Animals were allowed to recover on a warming mat.

### Mouse CNS Tissue Collection

Animals were euthanized 4 weeks after the ICM injection. Briefly, animals were deeply anesthetized with an intraparenchymal injection of Avertin (500 mg/kg; Sigma, St. Louis, MO). Mice were transcardially perfused via gravity perfusion with 20 mL of PBS (Cytiva; Marlborough, MA) supplemented with 100 U/mL heparin (Sandoz; Boucherville, QC, CA) over 20 min followed by 20 mL of 16% formaldehyde (Thermo Fisher, Waltham, MA) over 20 min. Brain, dorsal root ganglia and trigeminal ganglia were collected and stored overnight in formaldehyde at 4℃, then transferred to PBS and stored at 4℃.

### Mouse CNS Tissue Preparation

Tissue to be used for histological analysis were cryopreserved using an ascending sucrose gradient (15% sucrose, 0.02% sodium azide PBS solution for 24 h, followed by 30% sucrose, 0.02% sodium azide PBS solution until cryosectioning) at 4°C. Following cryopreservation, tissues were embedded in Tissue-Plus Optimal Cutting Temperature compound (OCT, Fisher Healthcare, Waltham, MA) and frozen in an ethanol-dry ice slurry. Embedded brains were sectioned sagittally at 40 µm on a cryostat. Sections were collected as free-floating slices in PBS supplemented with 0.02% sodium azide and stored at 4°C until use.

### Mouse CNS DAPI Staining and Slide Preparation

To visualize nuclei in brain, floating sections were placed in a 1:10,000 dilution of Hoechst nuclear dye (Invitrogen, Waltham, MA) for 10 min and washed for 5 min in PBS. Following staining, sections were mounted on SuperFrost Slides, with Prolong Gold Diamond mounting media (Invitrogen) and cover slipped. Mounted slides were cured at RT for 24h, then stored at 4°C until imaging.

### Mouse CNS Imaging

Confocal images were acquired using an inverted Nikon A1 confocal microscope equipped with 10× (Plan Apo, NA 0.45) and 20x objective lenses (Plan Apo NA 0.75) and a motorized stage. Three-color fluorescence imaging was conducted to visualize DAPI, FITC, and tdTomato signals. Specific excitation and emission filter settings were used to prevent signal overlap: DAPI (excitation: 405 nm, emission: 450/50 nm), FITC (excitation: 488 nm, emission: 525/50 nm), and tdTomato (excitation: 561 nm, emission: 595/50 nm). Images were acquired using the large image acquisition protocols within the NIS Elements software (Nikon Instruments, Lexington, MA). Acquisition parameters, including laser power, gain, and pinhole size, were standardized across samples.

### NHP CNS Injection via Intracisterna Magna (ICM)

The animal was started on a regimen of immunosuppressants (Prednisone 1 mg/kg; Cyclosporine A 8 mg/kg) daily, one week prior to the procedure. This regimen was continued daily through the day prior to necropsy. For the ICM injection, the animal was inducted with ketamine (5 mg/kg) and dexmedetomidine (0.02 mg/kg), then dosed with given maropitant citrate (1 mg/kg) and buprenorphine (0.2 mg/kg) and intubated. During procedure animal was maintained on isoflurane with oxygen inhalation to effect, 1-3%. A percutaneous ICM puncture with a 22-gauge Gertie Marx spinal needle was used to access the cisterna magna. Once CSF flow was observed, a manual bolus of 1 mL of an AAV pool (ATX002+wtAAV9) was injected over the course of a minute. Upon completion of dosing, the spinal needle was removed. Post-procedure, animal was provided atipamezole for anesthetic reversal and allowed to recover.

### NHP CNS Tissue Collection

The animal was euthanized 4 weeks after the ICM injection. Briefly, animals were sedated per NBR’s SOP and euthanized via an intravenous chemical overdose. The animal was perfused with chilled 0.001% sodium nitrite in saline, followed by 4% paraformaldehyde. A full gross examination was performed, and tissues (including whole brain, spinal cord, dorsal root ganglia, and peripheral organs) were collected and stored for 24-48 hours in paraformaldehyde at 4℃, then transferred to PBS and stored at 4℃.

### NHP CNS Tissue Preparation and Imaging

Paraformaldehyde-fixed sample was sectioned into 500 µm slabs using a megatome which were preserved with SHIELD reagents (LifeCanvas Technologies) using the manufacturer’s instructions (Park et al., 2018). Samples were delipidated for 14 days using LifeCanvas Technologies Clear+ delipidation reagents. Following delipidation samples were labeled using eFLASH (Yun et al., 2019) technology which integrates stochastic electrotransport (Kim et al. , 2015) and SWITCH (Murray et al., 2015), using a SmartBatch+ (or SmartLabel) device (LifeCanvas Technologies). Samples were incubated in 50% EasyIndex (RI = 1.52, LifeCanvas Technologies) overnight at 37°C followed by 1 d incubation in 100% EasyIndex for refractive index matching. After index matching, the samples were imaged using a MegaSPIM axially-swept light sheet microscope using a 1.6x objective (0.2 NA) with a 4 µm xyz voxel size (LifeCanvas Technologies). Three-color fluorescence imaging was conducted to visualize GFP (488 nm), tdTomato (561 nm) and autofluorescence (445 nm) signals.

### NHP CNS Analysis

Images were visualized using Imaris Explorer Software (Bitplane, Belfast, United Kingdom). Each region of interest was cropped and isolated using the BrainMaps: An Interactive Multiresolution Brain Atlas as a reference. Images were visually scored on a scale of 0 to ++++ indicating the amount of expression of each fluorophore.

### NHP CNS Tissue Preparation

Paraformaldehyde-fixed brain and spinal cord was sectioned into 10 mm slabs. These slabs were then cryopreserved using an ascending sucrose gradient (15% sucrose, 0.02% sodium azide PBS solution for 24 h, followed by 30% sucrose, 0.02% sodium azide PBS solution until cryosectioning) at 4°C. Following cryopreservation, tissues were embedded in Tissue-Plus Optimal Cutting Temperature compound (OCT, Fisher Healthcare, Waltham, MA) and frozen in an ethanol-dry ice slurry. Embedded slabs were sectioned coronally at 40 µm on a cryostat. Sections were collected as free-floating slices in PBS supplemented with 0.02% sodium azide and stored at 4°C until use.

### NHP CNS Staining and Slide Preparation

To reduce autofluorescence, floating sections were washed in TrueBlack® Lipofuscin Autofluorescence Quencher (#23007; Biotium) for 30 minutes at room temperature followed by a 30 second wash each in 70% ethanol and 50% ethanol, then a 5 min wash in wash buffer (0.3% Triton-X 100 in 1X PBS). Transgenes were counterstained as follows. Slices were incubated on a shaker at room temperature for 2 hours in blocking buffer (wash buffer with 5% NGS) followed by primary antibody in wash buffer shaking overnight at 4°C [chicken anti-GFP (1:6000; #GFP-1012; Aves Labs) and rabbit anti-RFP (1:2000; #600-401-379; Rockland)]. Slices were then washed three times for 5 minutes each in wash buffer at room temperature before incubation in secondary antibody in wash buffer for 2 hours shaking at room temperature [goat anti-chicken AlexaFluor 488 (1:1000; #A32931TR; Invitrogen) and goat anti-rabbit AlexaFluor 555 (1:1000; #A2148; Invitrogen)]. Slices were again washed three times for 5 minutes each in wash buffer at room temperature. To visualize nuclei in brain, floating sections were placed in a 1:1000 dilution in wash buffer of Hoechst nuclear dye (Invitrogen, Waltham, MA) for 20 minutes and washed three times for 5 minutes in wash buffer. To visualize neurons, floating sections were placed in a 1:50 dilution in wash buffer of NeuroTrace™ 640/660 Deep-Red Fluorescent Nissl Stain (ThermoFisher) for 1 hour and washed 3 times for 5 minutes in wash buffer. Following staining, sections were mounted on large glass slides (Fisher) with Prolong Diamond Antifade mounting media (Invitrogen) and cover slipped. Mounted slides were cured at RT for 24h, then stored at 4°C until imaging.

### NHP CNS Imaging

Confocal images were acquired using an inverted Nikon A1 confocal microscope equipped with a 10x (Plan Apo, NA 0.45) and 20x (Plan Apo, NA 0.8) objective lens and a motorized stage. Four-color fluorescence imaging was conducted to visualize DAPI, GFP, tdTomato, and Far-Red signals.

Specific excitation and emission filter settings were used to prevent signal overlap: DAPI (excitation: 405 nm, emission: 450/50 nm), GFP (excitation: 488 nm, emission: 525/50 nm), tdTomato (excitation: 561 nm, emission: 595/50 nm) and Far Red (exictation:640 nm, emission: 700/75 nm). Images were acquired using the large image acquisition protocols within the NIS Elements software (Nikon Instruments, Lexington, MA). Acquisition parameters, including laser power, gain, and pinhole size, were standardized across samples.

### NHP CNS Image Analysis

Images were visualized using Imaris Explorer Software (Bitplane, Belfast, United Kingdom). The Imaris Spots function was used to identify and count nuclei based on the DAPI channel and neurons based on the Nissl channel. Thresholding was based on an approximate size of 8 µm and 15 µm for nuclei and neurons, respectively. Positive cells in the GFP and tdTomato channels were also identified using the Imaris Spots function. ROIs were hand drawn with the Imaris Surface function based on the Nissl stain and used as a filter to identify DAPI+, Nissl+, GFP+ and tdTomato+ cells within these different ROIs. The DAPI+ spots were used to calculate the percent positive cells, and the Nissl+ spots were used to calculate the percent positive neurons.

### Molecular dynamics

Classical MD simulations of AAVs were performed to study the properties of loop inserts, both in the presence and absence of the AAV receptor (AAVR). Initial structures of single VP3 monomers for each variant were predicted using AlphaFold3 (AF3)^33^ with multiple sequence alignment to ensure topology close to the native AAV2 and previously reported 7m8 VP3s. These predicted VP3 structures were duplicated and aligned to the 7m8 cryo-EM structure VP3s (PDB: 6U0R) at the three-fold symmetry axis. Variant AAV-AAVR complexes were constructed by aligning our previously built 3-fold symmetric AAV variant models (in isolation) to the AAV2-AAVR cryo-EM complex structure (PDB: 6IHB) to get accurate positioning of the receptor to our new variant complexes.

Simulations were carried out using the pmemd module^34^ of GPU-accelerated Amber 22^35^ with the ff19SB force field^36^. The systems were solvated in TIP3P water^37^ within a periodic cuboid box with a 10 Å buffer from the protein surface. Water geometry was constrained using the SHAKE algorithm^38^, and the systems were neutralized with Na⁺ counterions. Long-range electrostatics were computed via the particle mesh Ewald method^39^ with 12 Å cutoffs for both electrostatics and Lennard-Jones interactions.

Energy minimization was performed for 30,000 steps, followed by gradual heating from 0 K to 300 K over 200 ps. Equilibration was then conducted under NPT conditions for 1 ns. Production simulations were run for 1 μs with a 2 fs timestep. No additional replicas were generated, as each system included 3 VP3s at the 3-fold axis, providing 3 replicate loop dynamics per variant.

Simulations were performed for atx002_ins_589, var16, AAV2, 7m8, and atx002, both in isolation and in complex with AAVR.

Post-simulation, clustering analysis was performed with CPPTRAJ^40^ using RMSD as the distance metric to identify the most populated conformations. H-bond analysis was also performed in CPPTRAJ using a 3.0 Å cutoff between donor heavy atoms and acceptors, and a minimum angle of 135° between the donor, hydrogen, and acceptor atoms. The cumulative contact frequency between residue pairs, calculated by summing H-bond contact fractions across all atom-atom pairs over the trajectory, indicates stronger contacts for residue pairs with more frequent or persistent interactions.

### Primers

Primer sequences are listed in Supplementary Table 2.

