## Supplemental materials for "Structural dynamics insights into principles underlying the fitness of new broadly potent AAVs"

### Supplementary figures

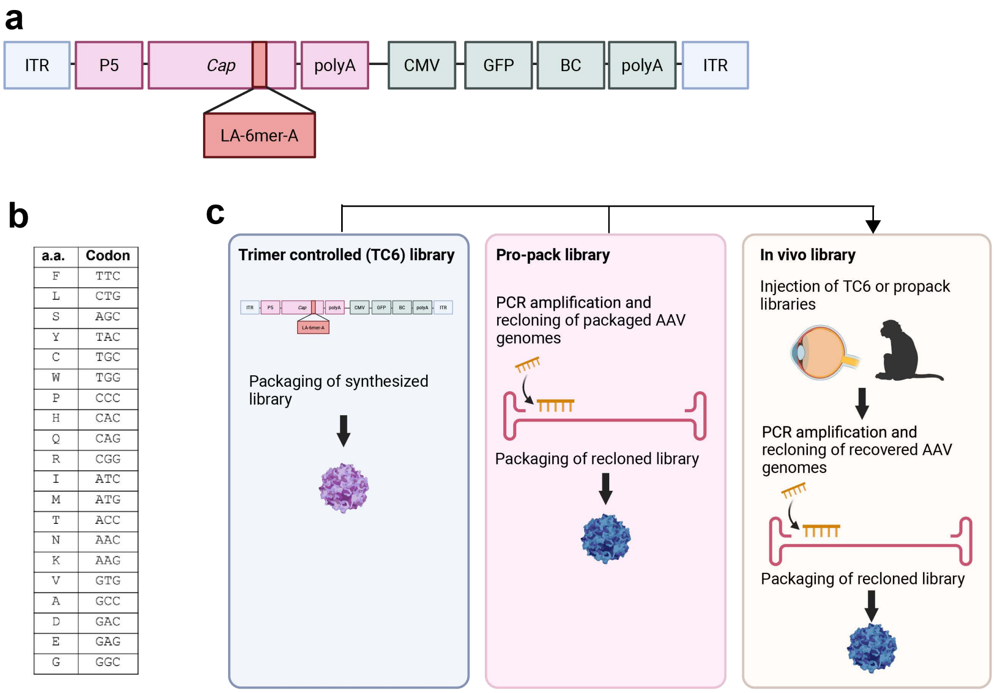

**Supplementary Figure 1: Library construction. a,** Map of the synthesized AAV library construct. **b,** Amino acid codons used in the trimer-controlled library **c,** Illustration showing the construction of the three libraries injected into NHPs. Trimer controlled libraries were packaged from synthesized DNA. Pro-pack library was created by PCR amplifying genomes from packaged AAVs in the TC6 library. The *in vivo* library was created by injecting TC6 or pro-pack libraries intravitreally into NHP eyes. AAV genomes were PCR amplified from retinal tissue, and then recloned into AAV backbones. All three libraries were injected into NHPs for scAAVengr-HUnT processing and analysis.

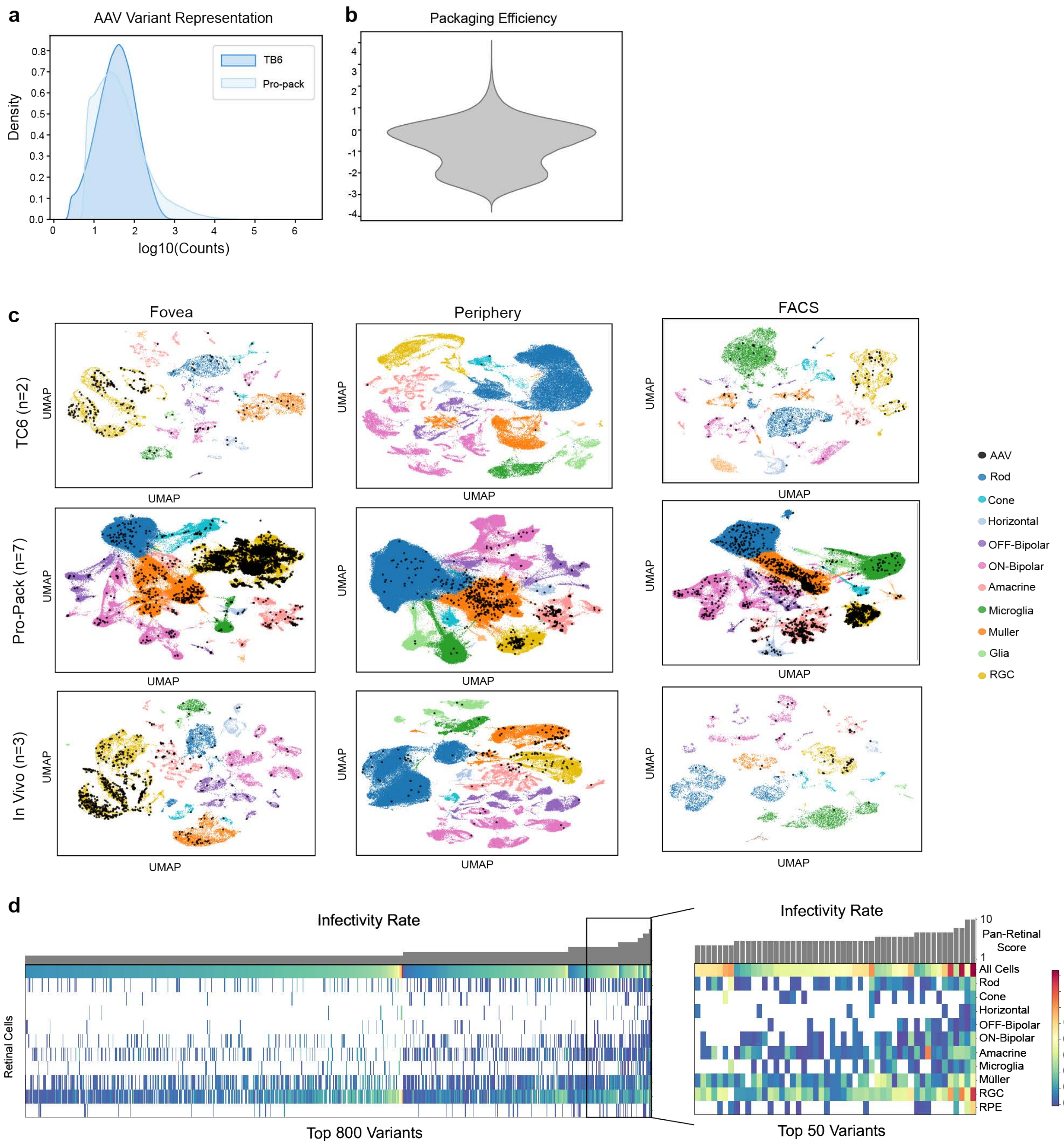

**Supplementary Figure 2: HT scAAVengr-HUnT variant analysis**. **a,** Representation of AAV variants in the TC6 and pro-pack libraries. **b,** Plot of AAV packaging efficiency derived from the TC6 library. Packaging Efficiency is calculated as follows: log10((pro-pack frequency/TC6 frequency])). **c,** UMAP plots of AAV variant transduction from the TC6, pro-pack and *in vivo* libraries. **d,** Heat map of AAV performance from the high-throughput round of scAAVengr screening.

a

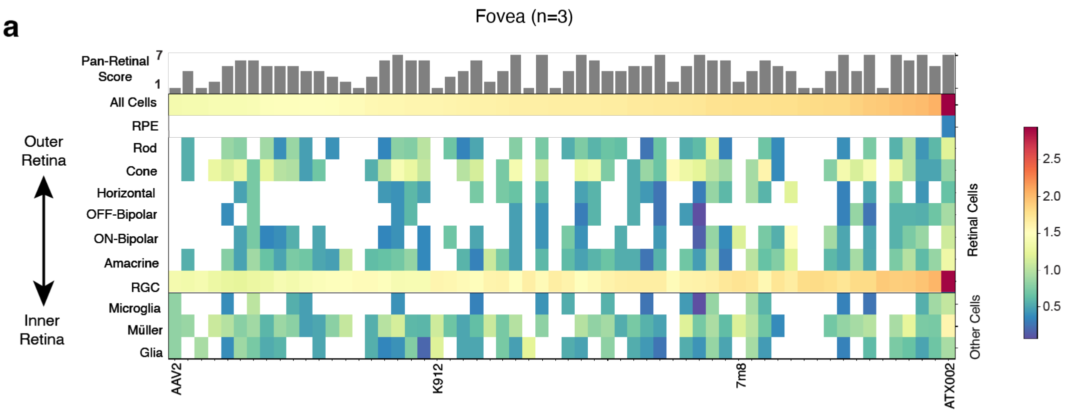

b

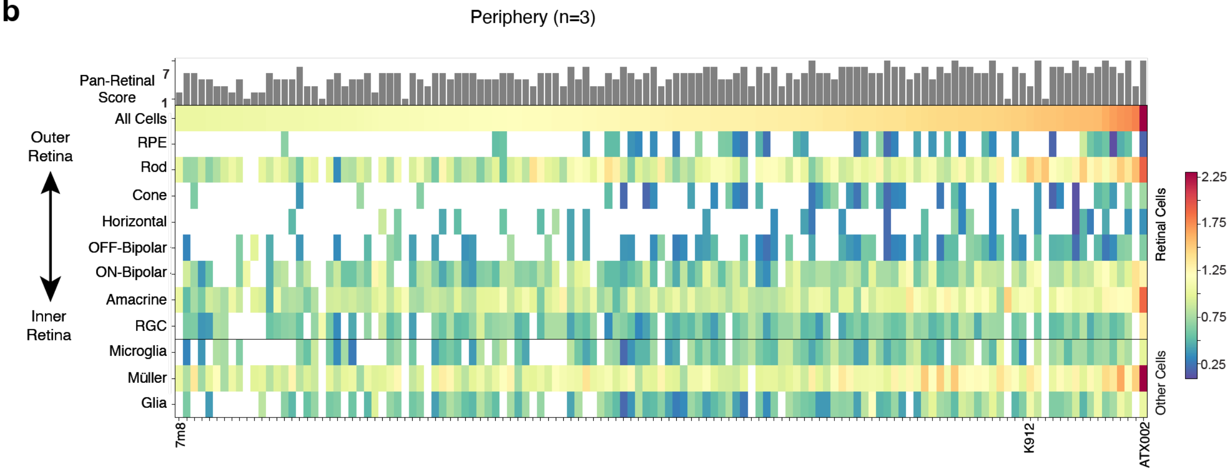

c

MAADGYLPDWLEDTLSEGIRQWWKLKPGPPPPKPAERHKDDSRGLVLPGYKYLGPFNGLDKG
EPVNEADAAALEHDKAYDRQLDSGDNPYLKYNHADAEFQERLKEDTSFGGNLGRAVFQAKKR
VLEPLGLVEEPVKTAPGKKRPVEHSPVEPDSSSGTGKAGQQPARKRLNFGQTGDADSVPDPQ
PLGQPPAAPSGLGTNTMATGSGAPMADNNEGADGVGNSSGNWHCDSTWMGDRVITTSTRT
WALPTYNNHLYKQISSQSGASNDNHYFGYSTPWGYFDFNRFHCHFSPRDWQRLINNNWGFR
PKRLNFKLFNIQVKEVTQNDGTTTIANNLTSTVQVFTDSEYQLPYVLGSAHQGCLPPFPADVFM
VPQYGYLTLNNGSQAVGRSSFYCLEYFPSQMLRTGNNFTFSYTFEDVPFHSSYAHSQSLDRLM
NPLIDQYLYYLSRTNTPSGTTTQSRLQFSQAGASDIRDQSRNWLPGPCYRQQRVSKTSADNN
NSEYSWTGATKYHLNGRDSLVNPGPAMASHKDDEEKFFPQSGVLIFGKQGSEKTNVDIEKVMI
TDEEEIRTTNPVATEQYGSVSTNLQRGN**LAEHQTRPA**RQAATADVNTQGVLPGMVWQDRDVY
LQGPIWAKIPHTDGHFHPSPLMGGFGLKHPPPQILIKNTPVPANPSTTFSAAKFASFITQYSTGQ
VSVEIEWELQKENSKRWNPEIQYTSNYNKSVNVDFTVDTNGVYSEPRPIGTRYLTRNL

**Supplementary Figure 3: Foveal and peripheral retinal heat maps from low-throughput round of scAAVengr-HUnT**. **a,** AAV performance in foveal punches. **b,** AAV performance in peripheral tissue. **c,** Sequence of ATX002 VP1.

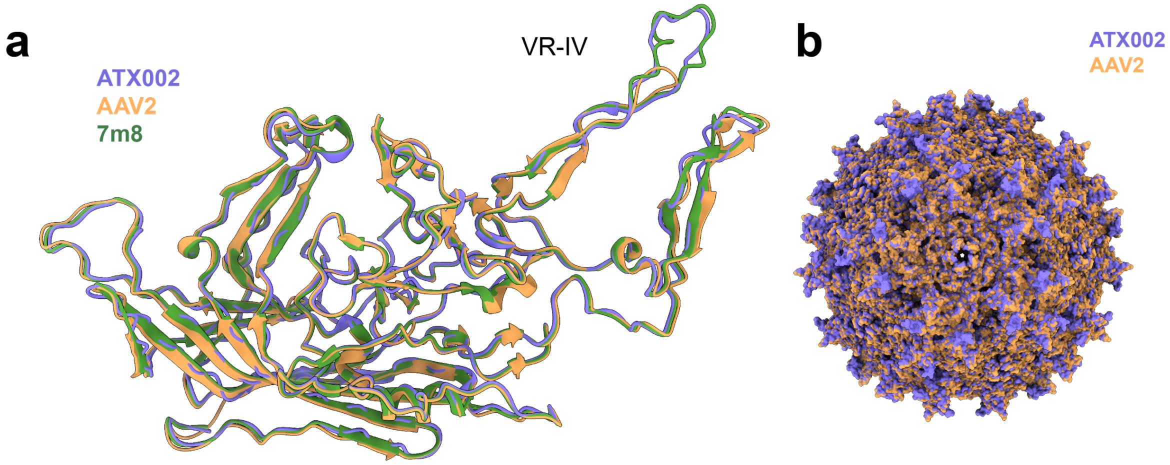

**Supplementary Figure 4: Overlay of ATX002, AAV2 and 7m8 structures.** ATX002 in purple, AAV2 in orange, and 7m8 in green. **a,** Overlay of ATX002, AAV2, and 7m8 VP3 subunits. **b,** Overlay of ATX002 and AAV2 capsid surfaces.

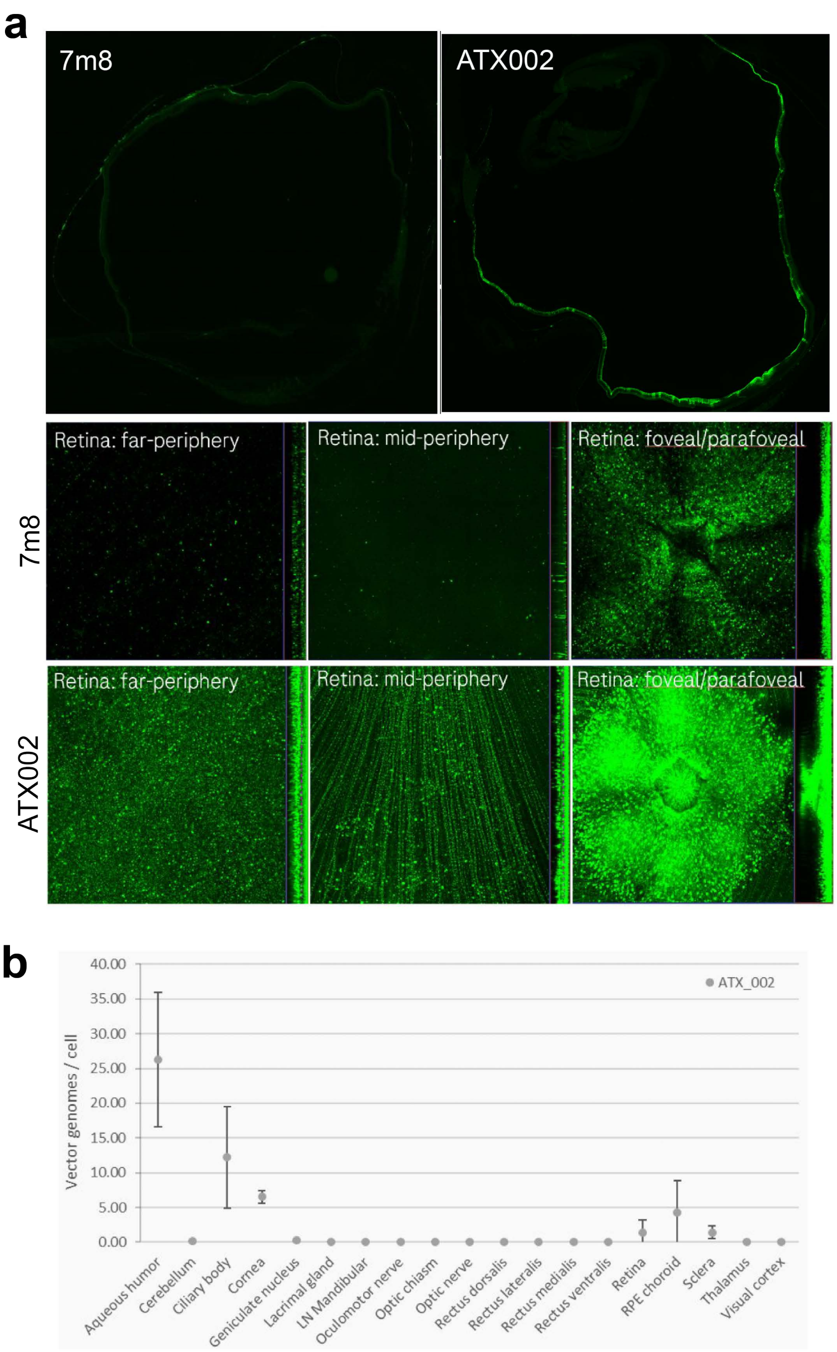

**Supplementary Figure 5: Improved performance of ATX002 in NHP retina. a,** GFP expression in whole eye cup sections (top row) far periphery, mid-periphery and fovea from NHPs injected with 7m8 and ATX002**. b,** Biodistribution of ATX002 following IVT injection in NHPs performed using multiplex ddPCR with RPP30 as the reference gene (n=3).

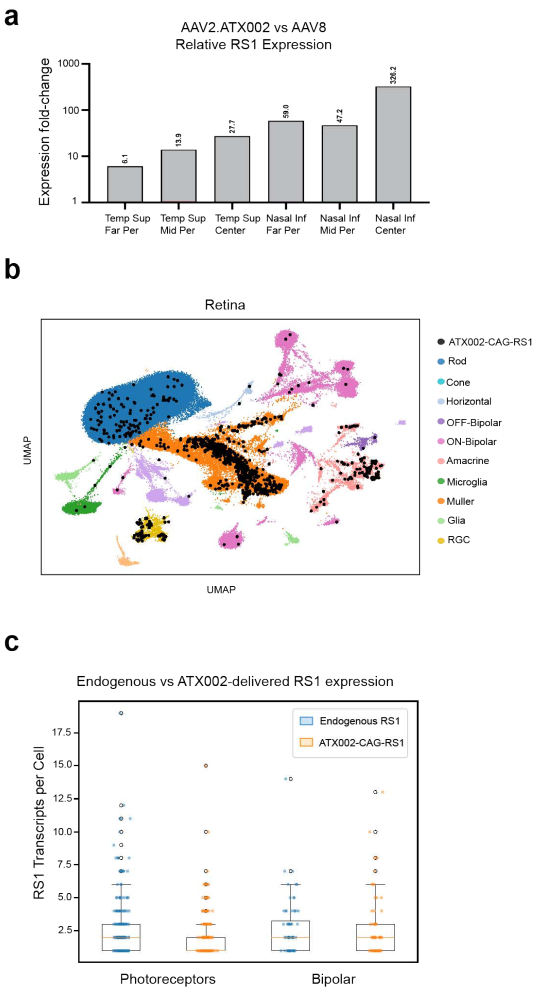

**Supplementary Figure 6: ATX002 delivery of human RS1 transgene in NHP via IVT injection.** X-linked retinoschisis (XLRS) is an early-onset inherited retinal dystrophy resulting from mutations in the *RS1* gene, which encodes a secreted protein involved in the maintenance of retinal cell architecture [14]. In clinical trials, IVT delivery of AAV-RS1 with wildtype AAV8 has been partially successful, although high doses that were associated with inflammation were required to achieve a partial response [16]. We therefore used ATX002 to deliver RS1 and compared performance to that of AAV8 following IVT injection in NHP (2E11 vg/eye, n=1 per serotype). **a,** Relative AAV-mediated RS1 expression compared to AAV8 across retinal regions. Quantitative RT-PCR showed that ATX002 outperformed AAV8 between 6.1-fold and 326-fold in relative RS1 transgene expression across peripheral regions of the retina. **b,** UMAP showing cell types transduced in ATX002-CAG-RS1-injected NHPs (n=2), dosed at a titer of 5E9 vg. Single cell analysis revealed RS1 expression in photoreceptors and bipolar cells, key cell types that are known to express RS1 and that are critical to target for gene augmentation therapy (fig. S6B) [15]. **c,** Levels of RS1 expression in transduced cells. Individual RS1 transgene transcripts were analyzed on a per-cell level and compared to endogenous RS1 expression. In transduced cells, transgene expression levels were similar to endogenous RS1 expression levels (fig. S6C).

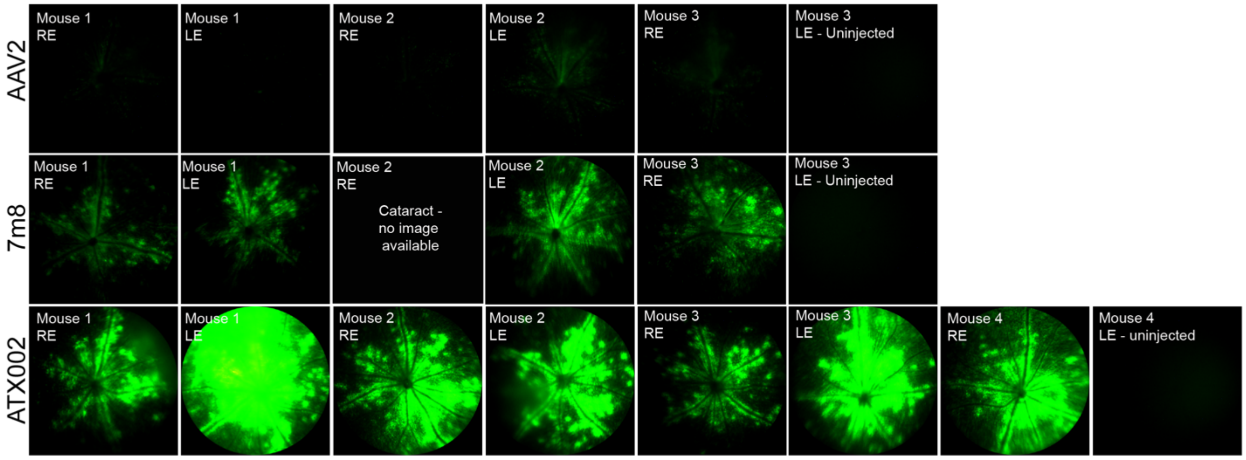

**Supplementary Figure 7: IVT injection of AAV2, 7m8 and ATX002 in mouse.** Fundus imaging from mice, injected bilaterally with AAV2, 7m8, or ATX002 and imaged one month after injection. In each group, one eye was left uninjected to evaluate potential contralateral spread. No GFP expression was observed in contralateral uninjected eyes.

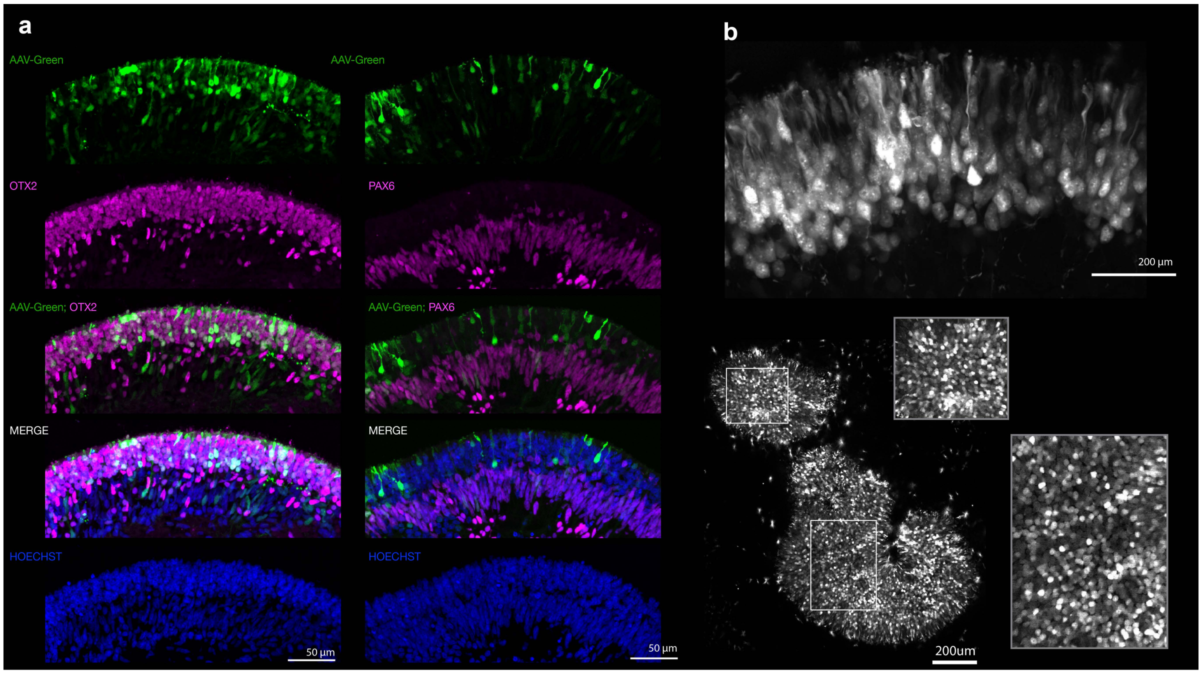

**Supplementary Figure 8: ATX002 expression in hiPSC-derived D135 human retinal organoids, 21 days post-infection. a,** Co-labeling of ATX002-CAG-mGL with OTX2 (photoreceptor and bipolar cell marker), PAX6 (bipolar and amacrine cell marker at this developmental stage). Hoechst labeling shows nuclei. **b, Top,** Optical section of an ATX002-treated D135 organoids imaged by two-photon microscopy. **Bottom**, max projection of optical sections focused on the outer surface of the organoid, revealing the mosaic of transduced photoreceptors (corresponding high-magnifications on the right).

.

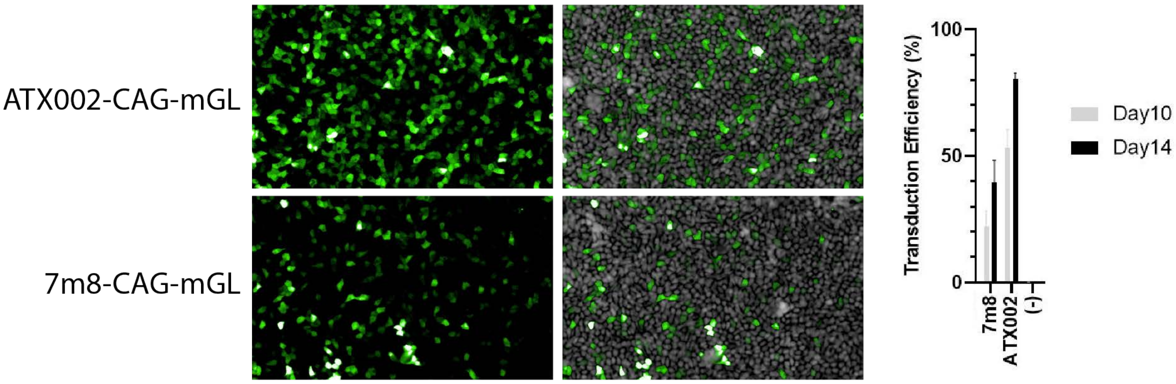

**Supplementary Figure 9: ATX002 transduces iRPE cells.** iCell RPE Cells were cultured and transduced with ATX002 or 7m8-CAG-mGreenLantern at MOI 2E4. Ten days and 2 weeks after AAV application, transduction efficiency was measured. ATX002 outperformed 7m8 at both timepoints.

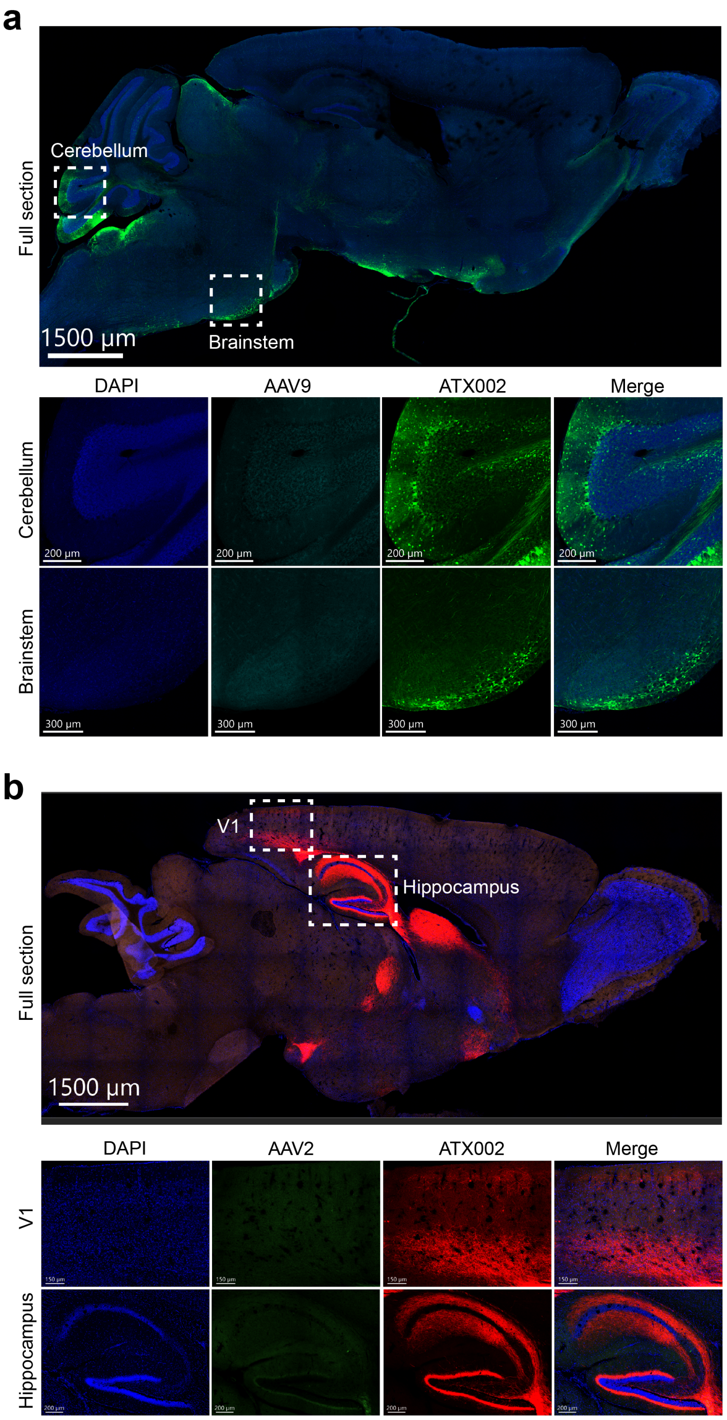

**Supplementary Figure 10: Mouse brain imaging. a,** ICM injections of AAV9 and ATX002 in mice. Imaging of cerebellum and brainstem show greater ATX002 transduction levels compared to AAV9. **b,** Intraparenchymal injections of AAV9 and ATX002 in mice. Imaging of V1 and hippocampus show dramatically greater ATX002 transduction levels compared to AAV2.

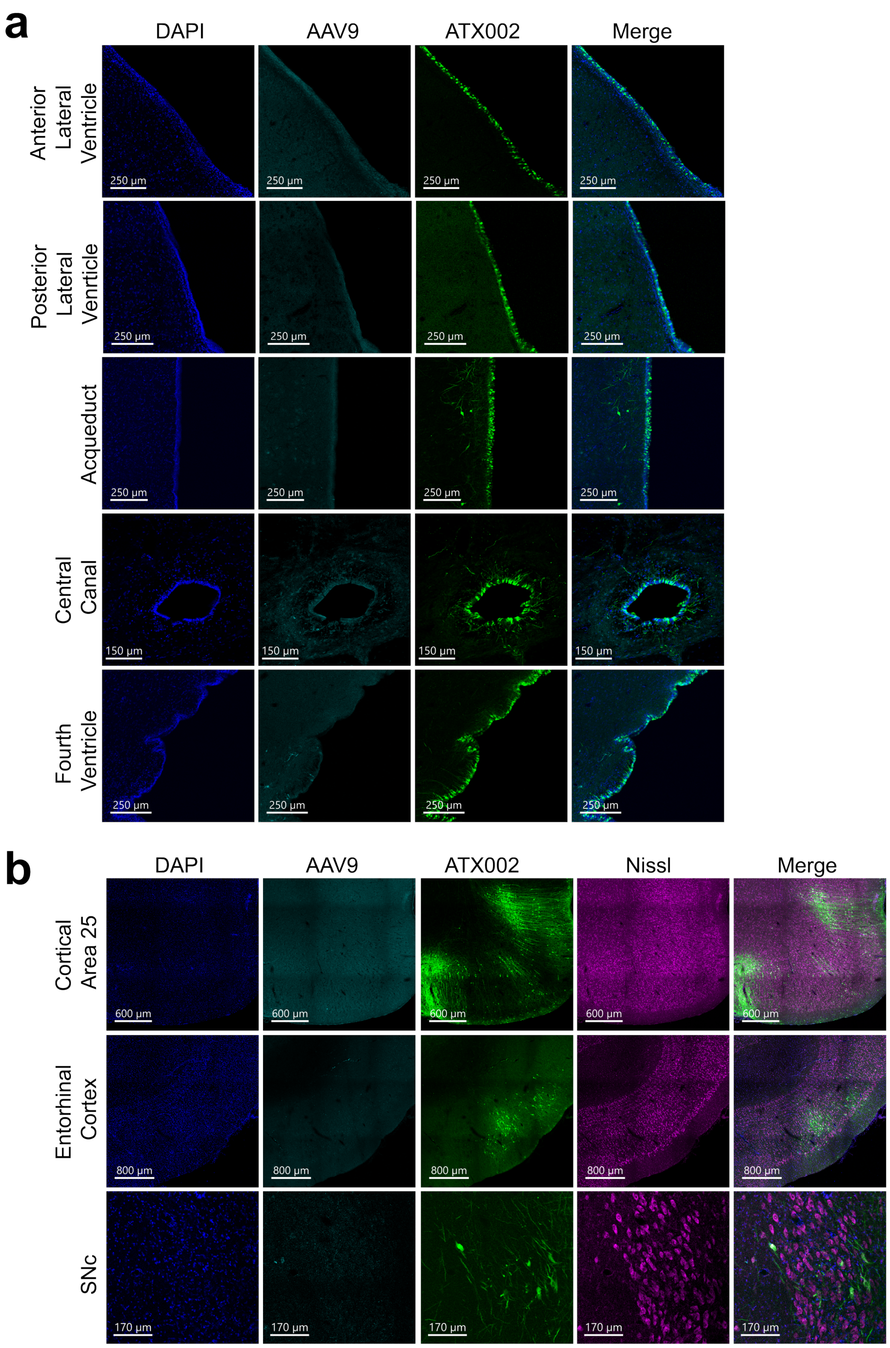

**Supplementary Figure 11: NHP brain imaging. a,** Imaging of ventricular system following ICM injections of AAV9 and ATX002 in NHP shows greater ATX002 transduction levels in ependymal cells compared to AAV9. **b,** Confocal imaging of brain regions show greater ATX002 transduction levels compared to AAV9. Colabeling with Nissl stain shows that transduction is largely neuronal.

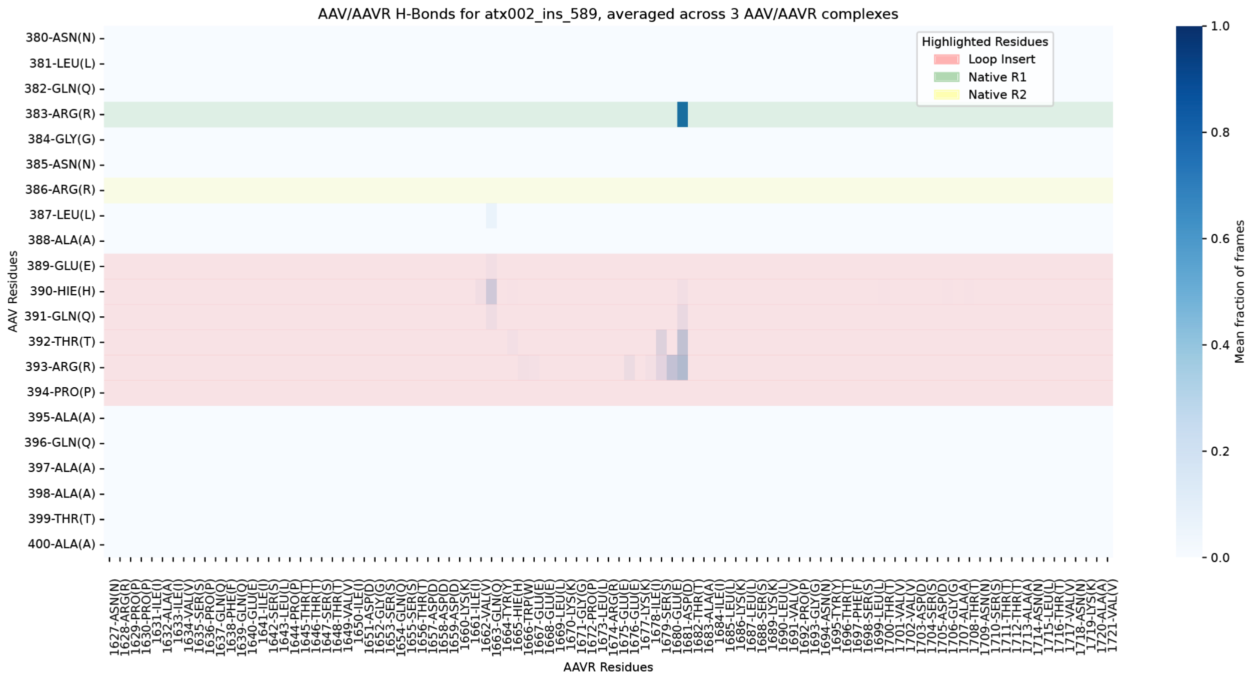

**Supplementary Figure 12:** AAV/AAVR H-Bonds for atx002_ins_589, averaged across 3 AAV/AAVR complexes.

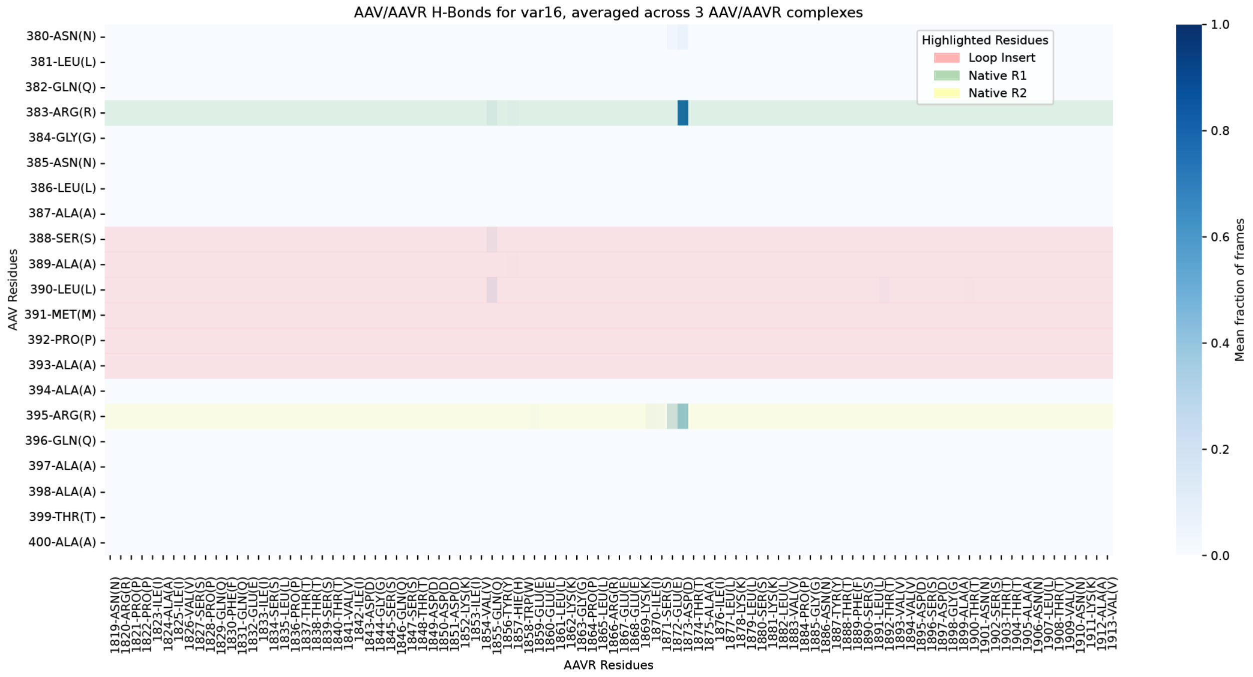

**Supplementary Figure 13:** AAV/AAVR H-Bonds for var16, averaged across 3 AAV/AAVR complexes.

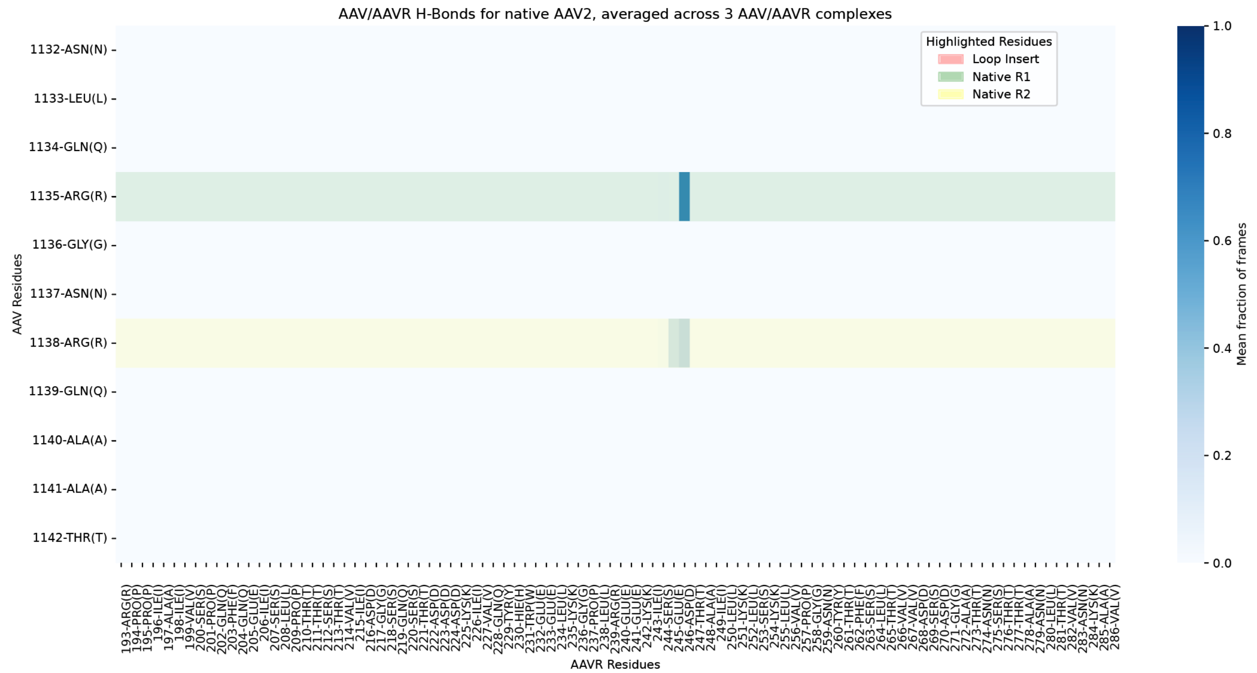

**Supplementary Figure 14:** AAV/AAVR H-Bonds for native AAV2, averaged across 3 AAV/AAVR complexes.

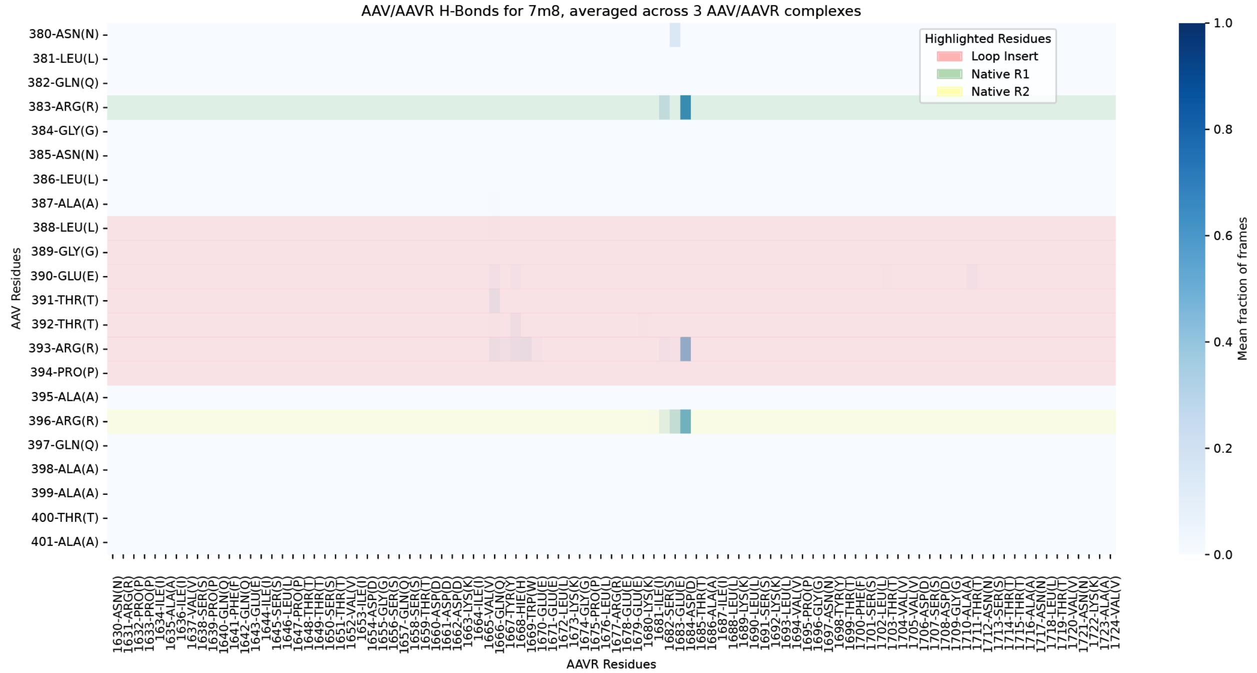

**Supplementary Figure 15:** AAV/AAVR H-Bonds for 7m8, averaged across 3 AAV/AAVR complexes.

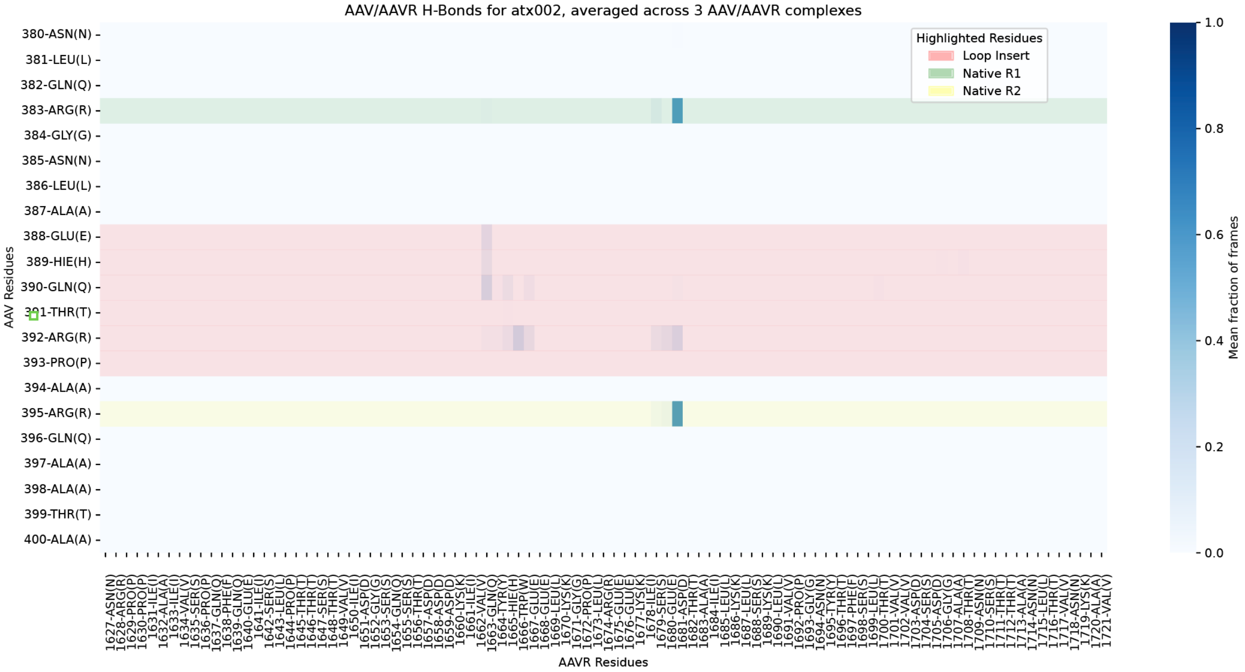

**Supplementary Figure 16:** AAV/AAVR H-Bonds for ATX002, averaged across 3 AAV/AAVR complexes.

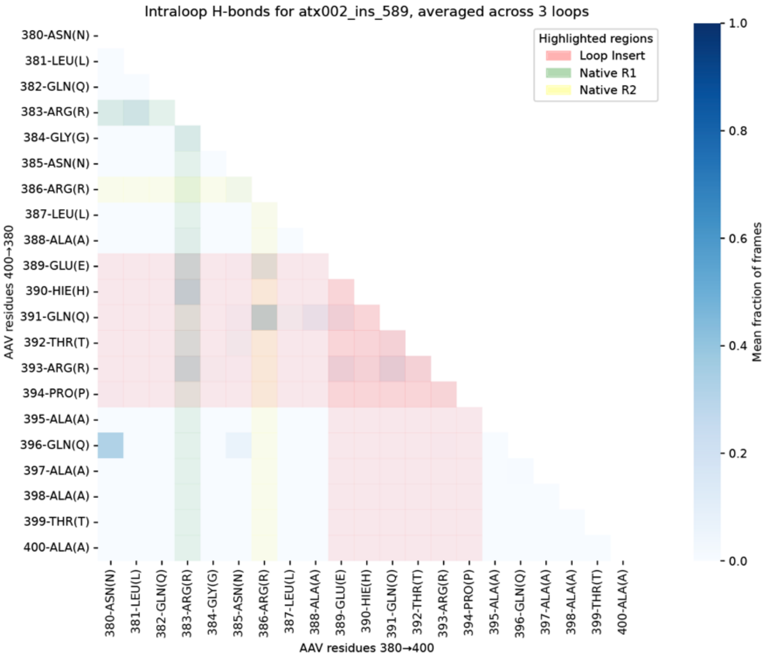

**Supplementary Figure 17:** Intraloop H-bonds for atx002_ins_589, averaged across 3 loops.

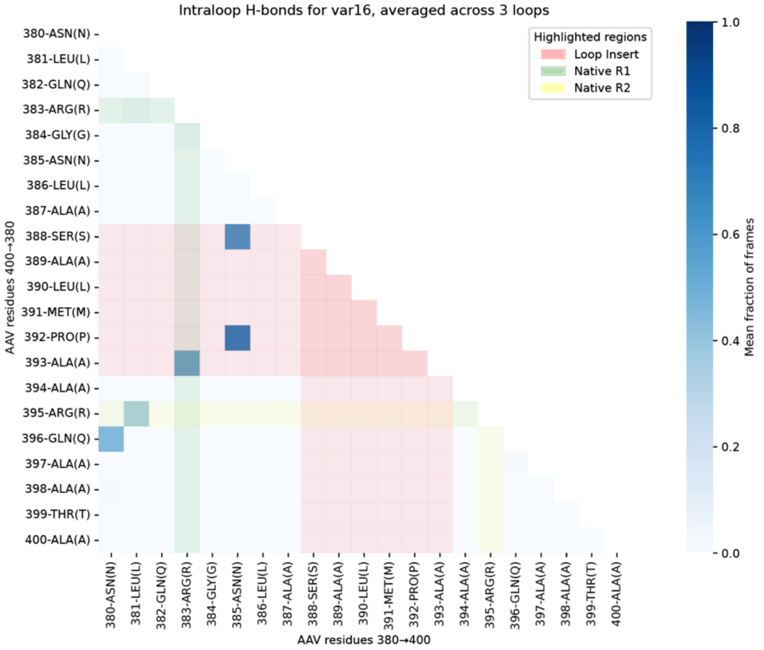

**Supplementary Figure 18:** Intraloop H-bonds for var16, averaged across 3 loops.

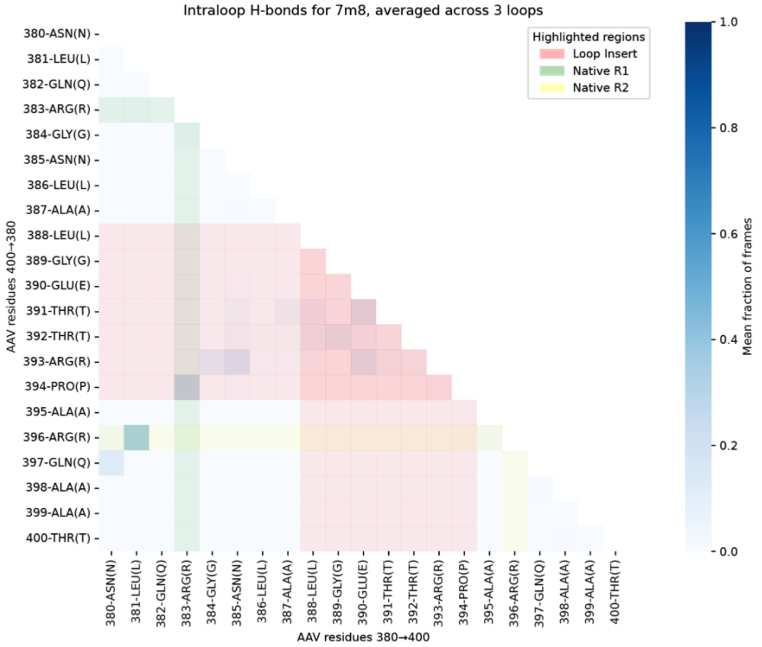

**Supplementary Figure 19:** Intraloop H-bonds for 7m8, averaged across 3 loops.

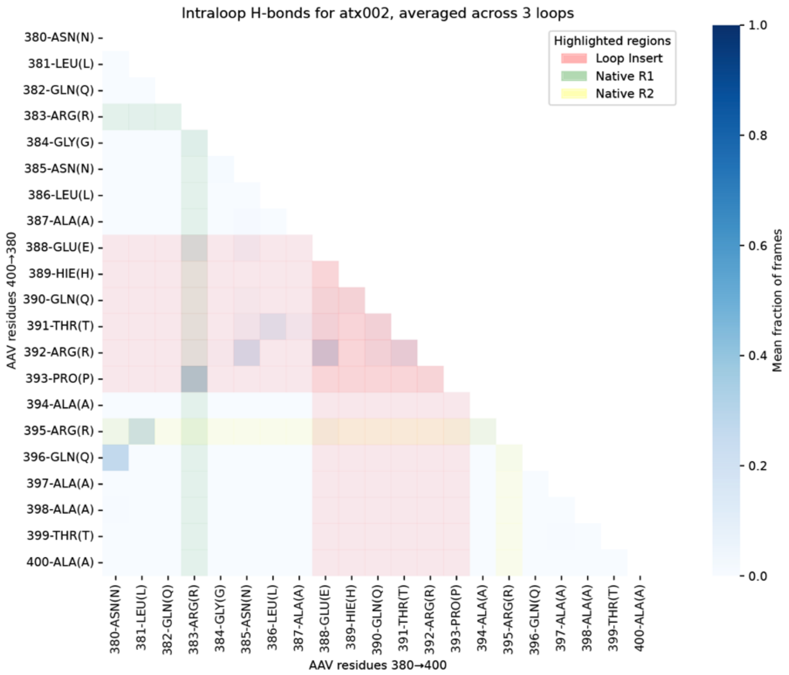

**Supplementary Figure 20:** Intraloop H-bonds for atx002, averaged across 3 loops.

### Supplementary tables

Supplementary Table 1. Summary of NHPs used in the studies.

| Species | DOB | M/F | Injection (left eye) | Injection (right eye) | NHP# | AAV2 Titer | Drugs administered | Adverse events |
|---|---|---|---|---|---|---|---|---|
| Rhesus macaque | 3/30/18 | M | Pro-pack library; 1.09E12 vg | TC6 library; 6.05E10 vg | M370-19 | 1:2 | Ceftriaxone (50 mg/kg 1x per day for 7 days, 2 rounds, metronidazole, 250 mg oral, 1x/day for 10 days) | Minor temporary periorbital swelling at time of injection |
| Cynomolgus macaque | 7/8/13 | F | Pro-pack library; 1.09E12 vg | TC6 library; 6.05E10 vg | M311-19 | 1:2 | Cyclosporine (oral, 7 mg/kg) once a day, starting 1 day after injection, Prednisolone and moxifloxicin eyedrops administered 2X daily starting 1dpi | Mild temporary uveitis |
| Cynomolgus macaque | 9/3/13 | F | Pro-pack library; 5.65E10 vg | TC6 library; 4.32E11 vg | M312-19 | 1:2 | Cyclosporine (oral, 7 mg/kg) once a day, starting 1 week before injection, 1x ketoprofen | Mild transient eye swelling |
| Cynomolgus macaque | 3/7/18 | F | TC6 library; 4.78E11 vg | N/a | M190-20 | 1:2 | Cyclosporine (oral, 7 mg/kg) once a day, starting 1 week before injection, ketoprofen 2 mg/kg x3 dpi | Mild temporary uveitis |
| Cynomolgus macaque | 8/14/18 | F | TC6 library; 4.78E11 vg | *In vivo* library; 1.87E11 vg | M193-20 | 1:2 | Cyclosporine (oral, 7 mg/kg) once a day, starting 1 week before injection, ketoprofen 2 mg/kg x3 dpi | Mild temporary uveitis |
| Rhesus macaque | 6/6/18 | F | TC6 library; 4.78E11 vg | *In vivo* library; 1.87E11 vg | M250-20 | 1:2 | Cyclosporine (oral, 7 mg/kg) once a day, starting 1 week before injection, ketoprofen 2 mg/kg x2 dpi | Mild temporary uveitis |
| Rhesus macaque | 4/21/18 | M | N/a | *In vivo* library; 1.87E11 vg | M251-20 | 1:2 | Cyclosporine (oral, 7 mg/kg) once a day, starting 1 week before injection | None |
| Rhesus macaque | 5/13/20 | F | N/a | Pro-pack library | M145-21 | 1:2 | Cyclosporine (oral, 8 mg/kg) and prednisone (oral, 1 mg/kg) once a day, starting 1 week before injection. 2mg/kg methylprednisone IM, 1x 1dpi. | None |
| Cynomolgus macaque | 2/25/10 | M | Pro-pack library | Pro-pack library | M286-21 | 1:2 | Cyclosporine (oral, 8 mg/kg) and prednisone (oral, 1 mg/kg) once a day, starting 1 week before injection. 2mg/kg methylprednisone IM, 1x 1dpi. | Uveitis in OS that reduced with methylprednisone. treatment |
| Cynomolgus macaque | 5/3/11 | M | Pro-pack library | Pro-pack library | M284-21 | 1:10 | Cyclosporine (oral, 8 mg/kg) and prednisone (oral, 1 mg/kg) once a day, starting 1 week before injection. 2mg/kg methylprednisone IM, 1x 1dpi. | Uveitis in OS, which resolved with methylprednisone treatment |
| Cynomolgus macaque | 12/12/13 | M | N/a | Pro-pack library | M289-21 | 1:10 | Cyclosporine (oral, 8 mg/kg) and prednisone (oral, 1 mg/kg) once a day, starting 2 weeks before injection. Oral ketoprofen 2x 1 and 2dpi. | None |

| | | | | | | | | |
|---|---|---|---|---|---|---|---|---|
| Cynomolgus macaque | 7/14/23 | M | Validation (200 variants); 5.25E11 vg | Validation (200 variants); 5.25E11 vg | M296-21 | 1:10 | Cyclosporine (oral, 8 mg/kg) and prednisone (oral, 1 mg/kg) once a day, starting 1 week before injection. 2mg/kg methylprednisone IM, 1x 1dpi. | Moderate uveitis, which resolved with methylprednisone. treatment |
| Cynomolgus macaque | 6/23/17 | M | Validation (200 variants); 5.25E11 vg | Validation (200 variants; 5.25E11 vg | M165-22 | 1:2 | Cyclosporine (oral, 8 mg/kg) and prednisone (oral, 1 mg/kg) once a day, starting 1 week before injection. 2mg/kg methylprednisone IM, 1x 1dpi. | Mild uveitis, which resolved with methylprednisone. treatment |
| Cynomolgus macaque | 2/23/16 | M | Validation (200 variants); 5.25E11 vg | Validation (200 variants); 5.25E11 vg | M29-23 | 1:10 | Cyclosporine (oral, 8 mg/kg) and prednisone (oral, 1 mg/kg) once a day, starting 1 week before injection. 2mg/kg methylprednisone IM, 1x 1dpi. | Mild uveitis, which resolved with methylprednisone. treatment |
| Cynomolgus macaque | 10/27/21 | M | ICM injection of ATX002-EF1a-tdTomato and AAV9-EF1a-eGFP; 5E12 vg of each vector | N/a | MB888A | Negative for AAV2 and AAV9 | Prednisone 1 mg/kg; Cyclosporine A, 8 mg/kg daily, starting one week prior to the procedure. Buprenorphine (0.2 mg/kg) at injection. | None |
| Cynomolgus macaque | 12/01/2020 | M | Validation AAV2.7m8-CAG-mGL 1E11 vg/eye | Validation AAV2.7m8-CAG-mGL 1E11 vg/eye | 2001 - 1035345 | Negative | 7mg/kg methylprednisolone IM, 1x 2 days before injection. Bilateral 100ul (40mg/mL) methylprednisolone subconjunctival, 1x 1dpi, 8dpi, 21dpi, 29dpi, 56dpi. | Moderate to severe uveitis manageable with methylprednisolone treatment |
| Cynomolgus macaque | 10/01/2020 | M | Validation AAV2.7m8-CAG-mGL 1E11 vg/eye | Validation AAV2.7m8-CAG-mGL 1E11 vg/eye | 2002 - 1035351 | Negative | 7mg/kg methylprednisolone IM, 1x 2 days before injection. Bilateral 100ul (40mg/mL) methylprednisolone subconjunctival, 1x 1dpi, 8dpi, 21dpi, 29dpi, 56dpi. | Moderate to severe uveitis manageable with methylprednisolone treatment |
| Cynomolgus macaque | 17/01/2020 | M | Validation AAV2.7m8-CAG-mGL 1E11 vg/eye | Validation AAV2.7m8-CAG-mGL 1E11 vg/eye | 2003 - 1035289 | Negative | 7mg/kg methylprednisolone IM, 1x 2 days before injection. Bilateral 100ul (40mg/mL) methylprednisolone subconjunctival, 1x 1dpi, 8dpi, 21dpi, 29dpi, 56dpi. | Moderate to severe uveitis manageable with methylprednisolone treatment |
| Cynomolgus macaque | 05/01/2020 | M | Validation ATX002-CAG-mGL 1E11 vg/eye | Validation ATX002-CAG-mGL 1E11 vg/eye | 3002 - 1035951 | Negative | 7mg/kg methylprednisolone IM, 1x 2 days before injection. Bilateral 100ul (40mg/mL) methylprednisolone subconjunctival, 1x 1dpi, 8dpi, 21dpi, 29dpi. | Moderate uveitis manageable with methylprednisolone treatment |
| Cynomolgus macaque | 23/01/2020 | M | Validation ATX002-CAG-mGL 1E11 vg/eye | Validation ATX002-CAG-mGL 1E11 vg/eye | 3003 - 1035991 | Negative | 7mg/kg methylprednisolone IM, 1x 2 days before injection. Bilateral 100ul (40mg/mL) methylprednisolone subconjunctival, 1x 1dpi, 8dpi, 21dpi, 29dpi, 56dpi. | Moderate uveitis manageable with methylprednisolone treatment |

| | | | | | | | | |
|---|---|---|---|---|---|---|---|---|
| Cynomolgus macaque | 2023 | M | N/a | Validation ATX002-CAG-mGL (1E+11 vg/eye) | M5-25 | Negative | Cyclosporine (oral, 8 mg/kg) and prednisone (oral, 1 mg/kg) once a day, starting 1 week before injection. | No inflammation noted |
| Cynomolgus macaque | Feb-3-2011 | M | N/a | ATX002-CAG-RS1 (5E+09 vg/eye) | M292-21 | 1:10 | Cyclosporine (oral, 8 mg/kg) and prednisone (oral, 1 mg/kg) once a day, starting 1 week before injection. | Moderate temporary uveitis |
| Cynomolgus macaque | Dec-12-2013 | M | ATX002-CAG-RS1 (5E+09 vg/eye) | N/a | M289-21 | 1:10 | Cyclosporine (oral, 8 mg/kg) and prednisone (oral, 1 mg/kg) once a day, starting 1 week before injection. | Moderate temporary uveitis |
| Cynomolgus macaque | Apr-21-2015 | M | ATX002-CAG-RS1myc (2.2E+11 vg/eye) | AAV8-CAG-RS1myc (2E+11 vg/eye) | M162-22 | 1:2 | Cyclosporine (oral, 8 mg/kg) starting 1 week before injection. Prednisone (oral, 1 mg/kg) once a day, starting on day of injection. | No inflammation noted |

Supplementary Table 2. List of primers used in the study.

| **Primer** | **Sequence** |
|---|---|
| F_1_Dict_Gib | AATGATACGGCGACCACCGAGATCTACACAGCGCTAGACACTCTTTCCCTACACGACGCTCTTCCGATCTTGGACGAGCTGTACAAGTAA |
| R_1_Dict_Gib | CAAGCAGAAGACGGCATACGAGATAACCGCGGGTGACTGGAGTTCAGACGTGTGCTCTTCCGATCTTGTGTTGACATCTGCGGTAG |
| Lib1_Gib_Dict_Insert_F | GAGGAAATCAGGACAACCAATCCCG |
| Lib1_Gib_Dict_insert_R | GGGGCAAACAACAGATGCTGGCAAC |
| Lib_amp_2_fwd_NheI | GCAGAGACTCTCTGGTGAATCCGGGCCCGGCCATG |
| ddPCR_target DNA_Fwd | AGGGTAGCCTGGAGAATTGC |
| ddCPR_target DNA_Probe | CCCAGCACTAGAAGTCGGCG |
| ddPCR_target DNA_Rev | TGATCACCGAATGGAAACAC |
| ddPCR_RPP30_Fwd | TGTGAACTTCAGGTGGCATTAC |
| ddPCR_RPP30_Probe | ACCCCAGTGTACCCAGGACAGT |
| ddPCR_RPP30_Rev | TCACAGAAACAGCAGGCATAAAC |
| RS1_Fwd_BC31 | CTAGCGAGCAGAAACTTATCTCCGAA |
| RS1_Rev_BC31 | GTGTACGACTATCACTCACATGCTA |
| RS1_Fwd_BC32 | GCGAGCAGAAACTTATCTCCGAAGA |
| RS1_Rev_BC32 | CGGTCGATGATGAGTGTACATGTGT |
| GAPDH_Fwd | ACAACAGCCTCAAGATCGTCAG |
| GAPDH_Rev | ACTGTGGTCATGAGTCCTTCC |
